# Inferring Telomerase Enzymatic Activity from Expression Data

**DOI:** 10.1101/2020.05.21.109249

**Authors:** Nighat Noureen, Shaofang Wu, Yingli Lyu, Juechen Yang, WK Alfred Yung, Jonathan Gelfond, Xiaojing Wang, Dimpy Koul, Andrew Ludlow, Siyuan Zheng

## Abstract

Active telomerase is essential for stem cells and most cancers to maintain telomeres. The enzymatic activity of telomerase is related but not equivalent to the expression of TERT, the catalytic subunit of the complex. Here we show that telomerase enzymatic activity can be robustly estimated from the expression of a 13-gene signature. We demonstrate the validity of the expression-based approach, named EXTEND, using cell lines, cancer samples, and non-neoplastic samples. When applied to over 9,000 tumors and single cells, we find a strong correlation between telomerase activity and cancer stemness. This correlation is largely driven by a small proliferating cancer cell population that exhibits both high telomerase activity and cancer stemness. This study establishes a novel computational framework for quantifying telomerase enzymatic activity and provides new insights into the relationships among telomerase, cancer proliferation, and stemness.

## INTRODUCTION

Telomerase is the ribonucleoprotein complex that adds telomeric repeats to telomeres at chromosome ends. In the absence of telomerase, telomeres progressively shorten due to incomplete replication of chromosome ends^1^. Persistent telomere shortening leads to senescence and crisis, thus continuously dividing cells including stem cells and most cancer cells require active telomerase to maintain telomere lengths^2^. Loss of telomerase activity results in degenerative defects and premature ageing^3,4^. In contrast, reactivation of telomerase enables malignant transformation and cancer cell immortality^5,6^. These observations emphasize the pivotal role of telomerase in many human health concerns and highlight the needs for close monitoring of its activity.

The telomerase complex is composed of the reverse transcriptase subunit TERT, the template containing non-coding RNA TERC, and accessory proteins such as dyskerin (DKC1) and telomerase Cajal body protein 1 (TCAB1). The core subunits are the catalytic subunit TERT and the RNA template TERC. In vivo, processive extension of telomeres, i.e. telomerase processivity, requires binding of telomerase to telomeres through a six-protein complex, shelterin, specifically its component protein TPP1^7,8^. Another shelterin protein POT1 bridges TPP1 and chromosome 3’ overhang, the DNA substrate of telomerase.

While *TERC* is thought to be abundant and ubiquitously expressed^9,10^, *TERT* is transcriptionally repressed in most somatic cells^11^. Some cancer lineages (e.g. brain, liver, skin and bladder) frequently acquire recurrent mutations in the *TERT* promoter region, predominantly at -124 and -146 loci upstream from *TERT* transcription start site^12–15^. These C>T mutations create consensus binding sites for GABP transcription factors, alter chromatin states, and enhance the transcriptional output of *TERT* ^16–20^. In bladder cancer, the promoter mutations correlate with increased *TERT* expression and telomerase enzymatic activity^21^.

Enzymatic activity is a fundamental metric of telomerase. Several protocols have been established to measure telomerase enzymatic activity, including the PCR-based TRAP assay and direct enzymatic assays^22–24^. These assays allowed for investigations of associations between telomerase activity and clinical and histopathologic variables in cancer and other diseases^25,26^, and regulation of telomerase activity by each component of the telomerase complex. Ectopic expression of *TERT* and *TERC*, in many cases *TERT* alone, increases telomerase enzymatic activity^27–29^. These data led to the view that *TERT* is the limiting component for telomerase activity.

However, new emerging data challenge the use of *TERT* expression as a surrogate for telomerase enzymatic activity. First, the *TERT* gene can transcribe more than 20 splicing isoforms^30^, but only the full-length transcript bearing all 16 exons can produce the catalytic subunit^31–33^. Secondly, single-cell imaging studies showed most *TERT* mRNAs localize in the nucleus but not the cytoplasm, and thus are not translated^34^.

Thirdly, endogenous TERT protein and *TERC* are far more abundant than the assembled telomerase complex in cancer cell lines^23^. Finally, *TERC* and accessory proteins can also impact telomerase activity. For example, telomerase activity in human T cells has been reported to relate to *TERC* levels rather than TERT^35^. In addition, mutations in *DKC1* and *TERC* both cause dyskeratosis congenita, a rare genetic syndrome related to impaired telomerase^10,36^.

Here we report a systematic analysis of telomerase activity in cancer. This analysis was enabled by a new telomerase activity prediction algorithm, <u>EX</u>pression-based <u>T</u>elomerase <u>EN</u>zymatic activity <u>D</u>etection (EXTEND).

## RESULTS

### Rationale and overview of EXTEND, a tool for predicting telomerase activity

An overview of the EXTEND algorithm is shown in **Supplementary Fig. 1**. We posited that comparing expression of telomerase positive and negative tumors could yield a telomerase activity signature. Most epithelial tumors express *TERT* and use telomerase to maintain telomeres. Without experimental evidence, however, *TERT* expression cannot reliably indicate positive telomerase for the reasons noted above. We instead used *TERT* promoter mutation to stratify such tumors, reasoning that the presence of this genetic change likely reflects evolutionary selection for telomerase as the predominant telomere maintenance mechanism (TMM). We further assumed that tumors with alternative lengthening of telomeres (ALT) phenotype, an alternate TMM mechanism, were negative controls. The ALT phenotype is usually determined through ALT-associated promyelocytic leukemia nuclear bodies, extrachromosomal telomeric DNA C-Circles, or ALT-associated telomere foci^37,38^. Large cohorts of experimentally confirmed ALT tumors are not yet publicly available, largely due to the rarity of the phenotype. However, mutations in *ATRX* and its interacting partner *DAXX* are nearly perfectly correlated with ALT ^39^. We thus searched the TCGA dataset for cancer types with both high frequencies of mutations in the *TERT* promoter and *ATRX* and *DAXX*. This search identified lower-grade glioma (LGG) (**Supplementary Fig. 2**), a cancer type demonstrating strong mutual exclusivity of the two TMMs^40^.

We identified 108 *TERT* co-expressing genes in the LGG dataset. These genes were further intersected with genes upregulated in the *TERT* promoter mutant tumors. The resulting 12 genes were complemented with *TERC*, the RNA subunit of the telomerase complex, giving rise to a 13-gene signature. Seven of the 13 genes were highly expressed in testis but low in other tissues (**Supplementary Table 1**). Mutations in *HELLS*, a gene encoding a lymphoid-specific helicase, cause the centromeric instability and facial anomalies (ICF) syndrome, a genetic disorder associated with short telomeres^41^. Functional enrichment analysis suggested an overrepresentation of the signature genes in the cell cycle pathway (FDR=1.95e-4), particularly S phase (FDR=0.01, **Supplementary Table 2**), a narrow time window when telomerase is active in extending telomeres^42^.

To score this signature, we designed an iterative rank-sum method (**Supplementary Fig. 1**). This method first divides the signature into a constituent component (*TERT* and *TERC*) and a marker component (the other 11 genes). The constituent component is scored by the maximum ranking of *TERT* and *TERC*, whereas the marker component is scored by the rank sum of the signature genes. Because the size of the constituent component is much smaller than the marker component, we adjusted its contribution to the final score by a factor determined based on the correlation between the original constituent component score and the aggregated score. Using cancer cell lines from the Cancer Cell Line Encyclopedia (CCLE), we estimated that the constituent component generally contributes less than 20% to the final scores (**Supplementary Fig. 3**).

Since *TERC* lacks a long poly(A) tail, its measurement is less robust with a poly(A) enriched mRNA sequencing protocol. Thus, we tested the stability of EXTEND across sequencing protocols. Using samples sequenced by both ribosomal RNA depletion and poly(A) enrichment protocols^43^, we found *TERC* was indeed less concordant than *TERT* between the two protocols (Rho=0.71 vs 0.98), but EXTEND scores nevertheless agreed well (Rho=0.96, P=1.9e-6; **Supplementary Fig. 4**).

### Validation and comparison with *TERT* expression in cancer

We first validated EXTEND with cancer cell lines. We performed semi-quantitative TRAP assays on 28 patient-derived glioma sphere-forming cells (GSCs) (Methods). EXTEND scores were significantly correlated with experimental readouts (Rho=0.48, P=0.01; **Fig. 1A**). Although this correlation was comparable to *TERT* expression (Rho=0.5, P=0.01), EXTEND demonstrated superior performance in differentiating two recently determined ALT lines, GSC5-22 and GSC8-18^37^ (**Supplementary Fig. 5**). Using 11 bladder cancer lines with available RNAseq data, we compared predictions obtained using EXTEND with more accurate direct enzymatic assays^21^. EXTEND scores and cell line telomerase activity were significantly correlated (Rho=0.72, P=0.01), whereas the correlation for *TERT* did not reach statistical significance (Rho=0.55, P=0.08) (**Fig. 1B, Supplementary Fig. 6**). We then measured telomerase activity of 15 lung cancer cell lines using digital droplet TRAP assays^24^. EXTEND outperformed *TERT* expression in predicting telomerase on these cell lines (Rho=0.65, P=0.01 vs Rho=0.48, P=0.07) (**Fig. 1C**, **Supplementary Fig. 7**).

**Fig. 1.**
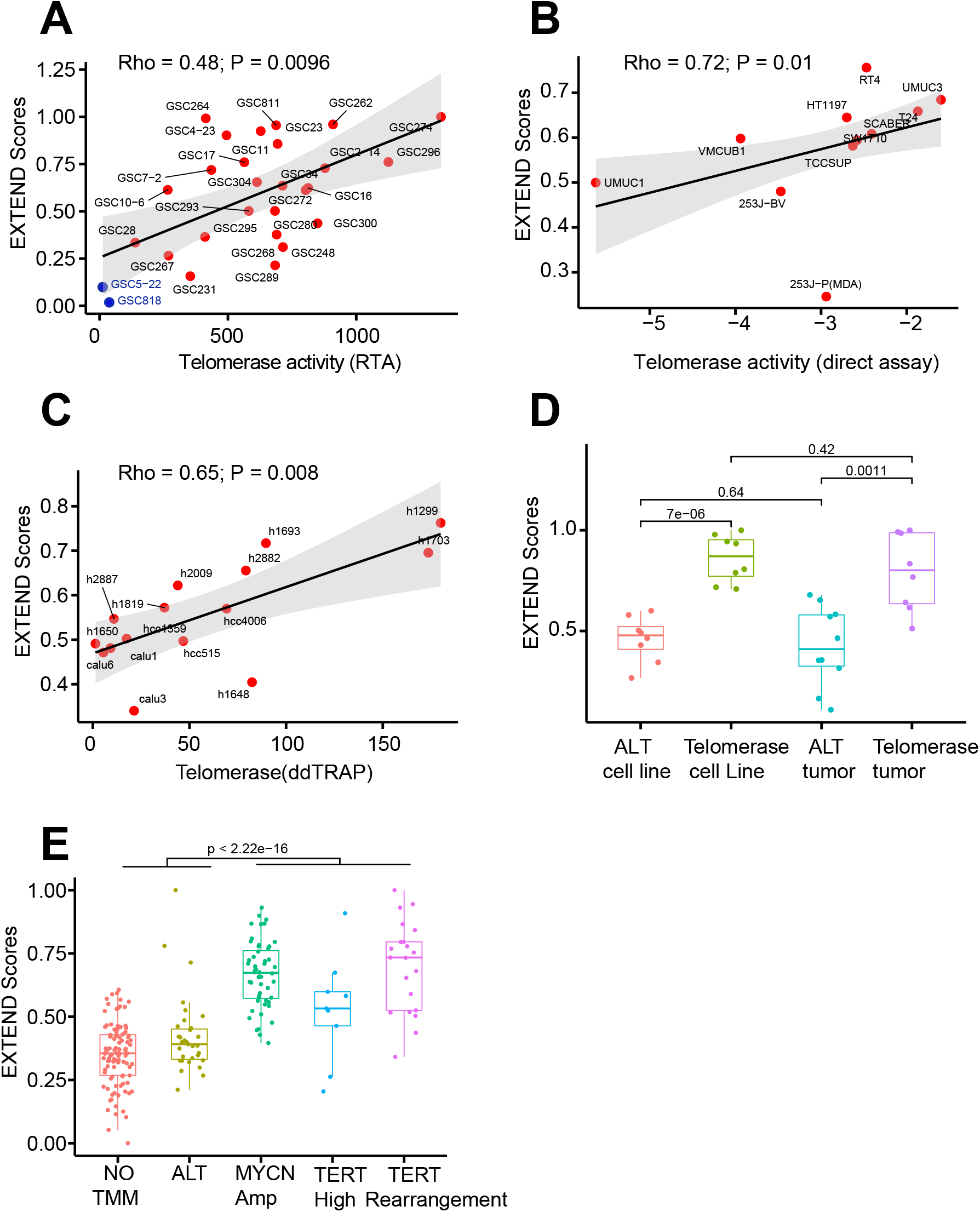
EXTEND Validation. **(A)** Correlation between EXTEND score and TRAP assay readouts in 28 glioma sphere forming cell lines. Two ALT cell lines are labeled in blue. Spearman correlation was used to calculate p value and Rho for (A-C). **(B)** Correlation between EXTEND score and direct enzymatic assay results in 11 bladder cancer cell lines. **(C)** Correlation between EXTEND score and digital droplet TRAP assay in 15 lung cancer cell lines. **(D)** EXTEND scores across ALT and telomerase positive tumors in Liposarcomas (GSE14533). Both telomerase positive tumors and cell lines show significantly higher EXTEND scores than ALT samples. **(E)** EXTEND scores across the five TMM groups of Neuroblastoma (GSE120572). Telomerase positive groups (MYCN amplification, TERT high and TERT rearrangement) show significantly higher scores than ALT and no-TMM groups.

We next tested EXTEND in two cancer types enriched with ALTs^44^, liposarcoma and neuroblastoma. In liposarcoma^45^, telomerase-positive tumors had significantly higher EXTEND scores than ALT tumors (P<0.01, Student t test; **Fig. 1D**). Neuroblastomas were previously divided into five groups based on telomere maintenance mechanisms: *TERT* promoter rearrangement, *MYCN* amplification, *TERT* expression high, ALT, and no TMM^38^. Enzymatic assays suggested that the first three groups were telomerase-positive, whereas the latter two had very low telomerase activity^38^. Consistent with these observations, EXTEND estimated higher telomerase activity in the three telomerase-positive groups (**Fig. 1E**). The *TERT* high group was estimated to have relatively lower scores than the other two telomerase-positive groups, a pattern also consistent with the experimental data^38^.

### Telomerase activity in non-neoplastic and embryonic samples

We then applied EXTEND to tissue samples from the GTEx dataset (**Supplementary Table 3**). These samples were collected post-mortem from healthy donors, except for a few transformed cell lines. Among the 52 sub-tissue types, we observed the highest EXTEND scores in Epstein-Barr virus-transformed lymphocytes, testis, transformed skin fibroblasts, and esophageal mucosa, a tissue with a high self-renewal rate (**Fig. 2A, Supplementary Fig. 8**). The testis was recently shown to have the longest average telomere lengths among human tissues^46^. Highly differentiated tissue skeletal muscle samples had the lowest average scores. Skin-transformed fibroblasts had almost negligible *TERT* expression despite a relatively high EXTEND score. In contrast, brain tissues, including substantia nigra (brain_SN), putamen, nucleus accumbens (brain_NA) and caudate, had low EXTEND scores but expressed *TERT* at a detectable level. To explain this disparity, we examined *TERT* alternative splicing. Compared with testis, most expressed *TERT* were short-spliced forms in the brain thus could not encode catalytically active proteins (**Supplementary Fig. 9**).

**Fig. 2.**
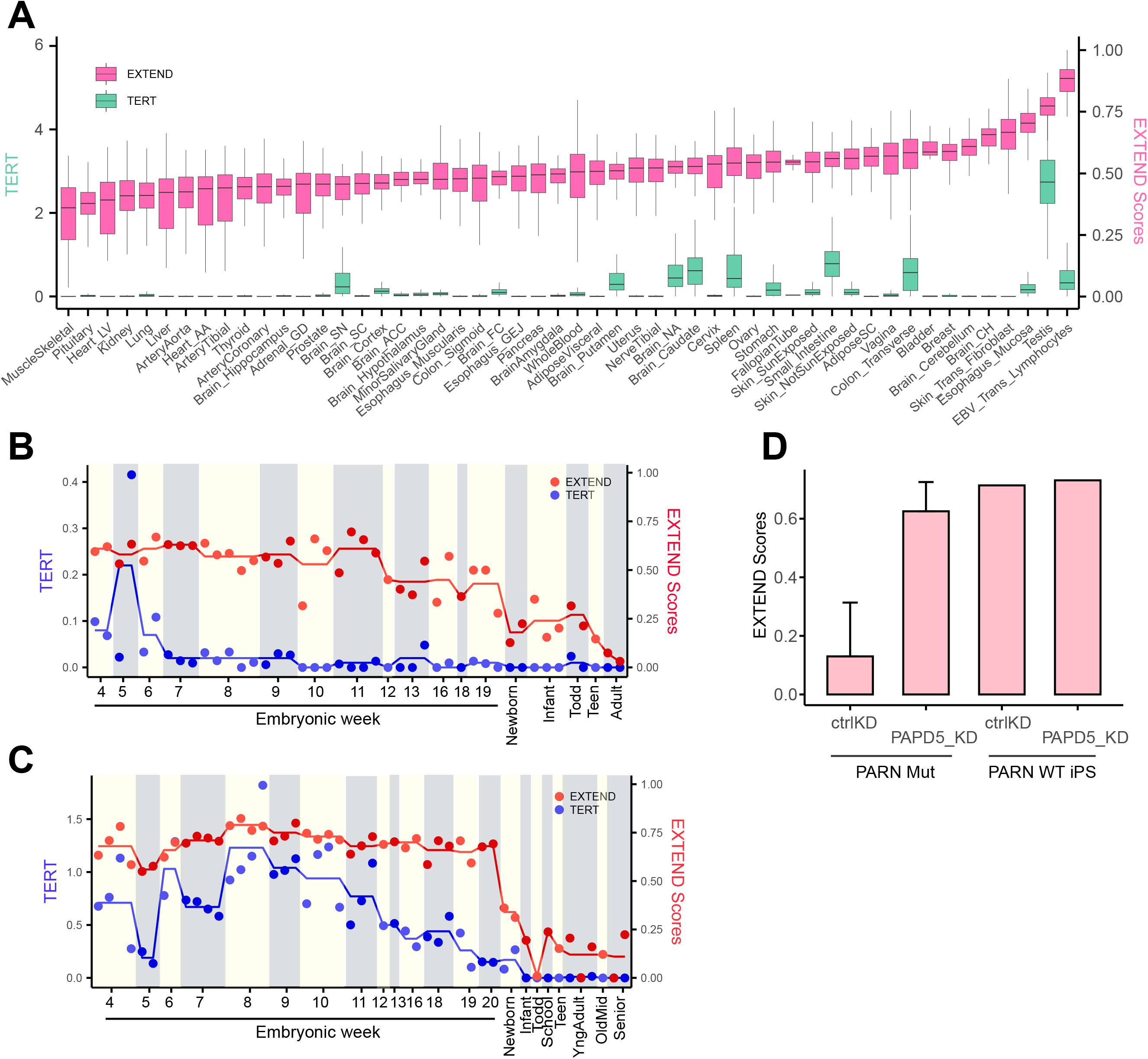
EXTEND scores across normal tissues and during embryonic development. **(A)** EXTEND scores and TERT expression across 52 sub tissues across Genotype-Tissue Expression (GTEx) data. The left y-axis indicates TERT expression while the right y-axis represents EXTEND scores. **(B and C)** TERT expression (blue) and EXTEND scores (red) across tissue development phases in (**B**) heart tissue and (**C**) liver tissue. Red and blue lines represent the mean of EXTEND scores and TERT expression of samples from each age group. **(D)** EXTEND predicts higher telomerase activity in PARN mutant, PARD5 knockout sample (left two bars), whereas no such effect is observed in PARN wildtype iPS cells (right two bars). Data is downloaded from GSE81507.

Next, we analyzed human tissues during embryonic development. We focused on heart and liver, two organs with distinct self-renewal behaviors. Cardiomyocytes were thought to lose their proliferative ability after birth^47^, whereas telomerase-expressing hepatocytes robustly repopulate the liver in homeostasis throughout adulthood^48,49^. Using a recently published dataset^50^, we analyzed both organs through embryonic weeks 4-20 and after birth. For heart, EXTEND scores dropped considerably between the 12^th^ and 13^th^ embryonic weeks but remained at modest levels in infants and toddlers (**Fig. 2B**). A further drop was observed when entering adulthood. These observations agreed with previous studies that reported decreased telomerase activity after the 12^th^ embryonic week in the heart tissue^51^, and that infant hearts have higher telomerase activity than those of adults^52^.

We did not observe significant decreases in EXTEND scores for liver during late embryonic weeks, despite the declining *TERT* expression after around 10 weeks. EXTEND scores largely remained stable across the lifespan, although their levels were much lower than in fetal samples (**Fig. 2C**, P=1.1e-9).

Finally, we applied EXTEND to samples from a patient with dyskeratosis congenita^53^. This patient carried loss-of-function mutations in Poly(A)-specific ribonuclease (*PARN*), a gene required for *TERC* maturation. Inhibition of poly(A) polymerase PAP-associated domain–containing 5 (*PAPD5*) counteracts *PARN* mutations to increase *TERC* levels and telomerase activity ^53^. EXTEND predicted higher telomerase activity in *PARN* mutant cells compared with wild-type controls upon *PAPD5* knockdown, confirming the earlier finding (**Fig. 2D**).

### Landscape of telomerase activity in cancer

We next analyzed more than 9,000 tumor samples and 700 normal samples from TCGA (**Supplementary Table 4**). We first evaluated if EXTEND scores were also reflective of telomerase processivity in cancer. POT1 is a shelterin component that regulates telomerase processivity through interactions with chromosome 3’ overhang and TPP1, the telomerase anchor to telomeres^54^. TPP1-POT1 heterodimer has been reported to enhance telomerase processivity^55–57^. EXTEND scores were positively correlated with *POT1* expression across pan-cancer (R=0.22, P<2.2e-16). No correlation was observed between *TERT* and *POT1* (R=-0.004, p=0.71) (**Supplementary Fig. 10**). We did not find significant associations between telomere length and EXTEND scores in TCGA cohorts after adjusting for multiple hypothesis testing (**Supplementary Fig. 11**).

As expected, tumors had significantly higher EXTEND scores than matched normal samples (**Fig. 3A**; P < 2.2e-16, t test). Similar to the GTEx sample analyses, scores varied across normal tissues. Gastrointestinal organs (esophageal – ESCA, stomach – STAD, colorectal - CRC), mammary gland (breast - BRCA), and reproductive organs (uterus endometrial - UCEC) had overall higher scores. EXTEND scores varied across cancer types, with kidney (KIRP, KIRC), thyroid (THCA), prostate (PRAD), and pancreatic (PAAD) demonstrating the lowest scores (**Fig. 3A**). These cancer types also appeared to have the smallest differences between normal and cancer samples. The small difference for pancreatic cancer may reflect this cancer type’s high impurity.

**Fig. 3.**
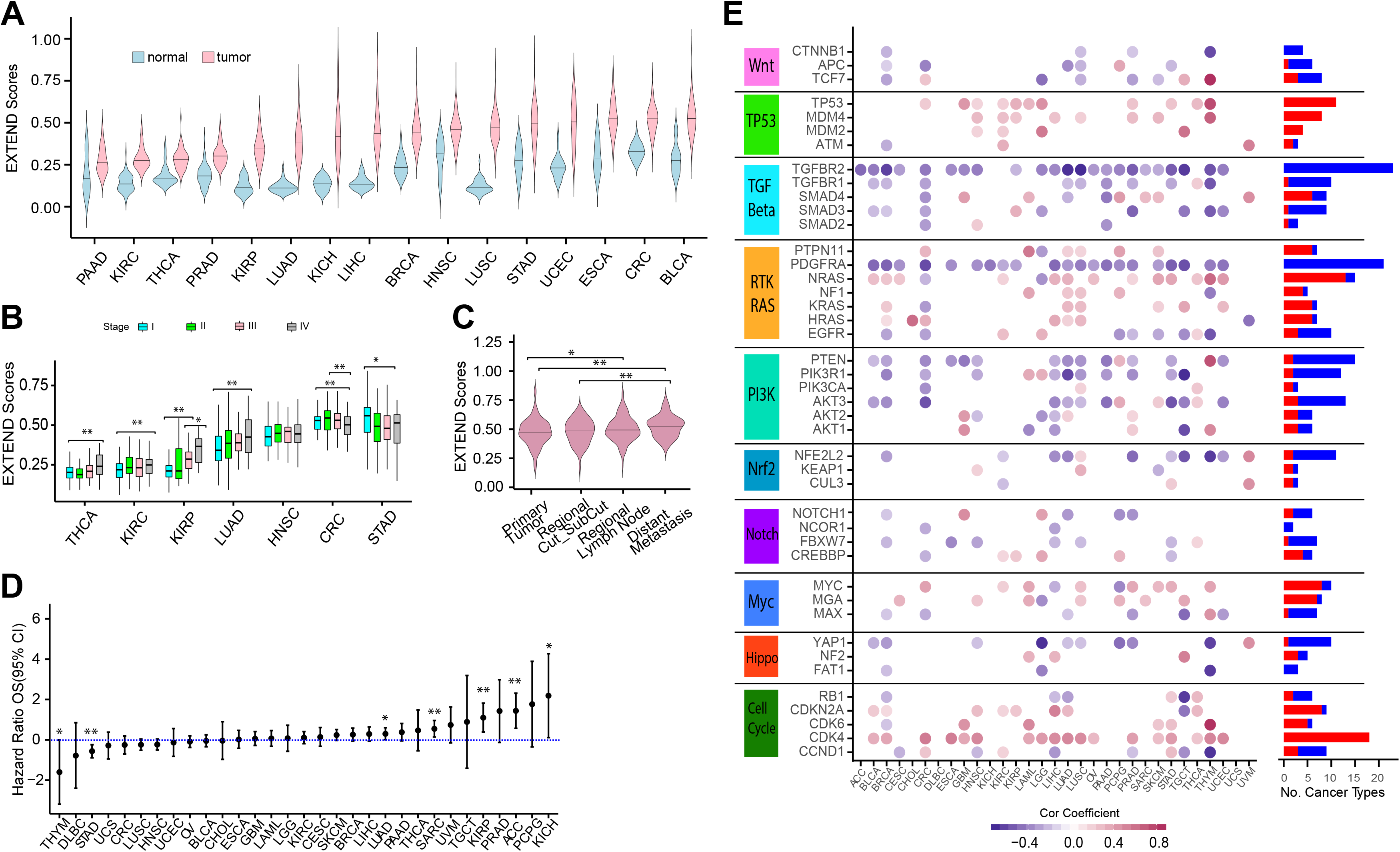
EXTEND scores in cancer. **(A)** EXTEND scores for TERT-expressing tumors and normal samples across 16 TCGA cancer cohorts. **(B)** EXTEND scores across tumor stages. Only tumor types with a minimum of 10 cases for each stage were used in this analysis. P-values were calculated using two-sided t-test. *, P<0.05; **, P< 0.01. **(C)** EXTEND scores of cutaneous melanoma (TCGA SKCM). **(D)** Hazard ratio plot based on univariate cox regression model for 31 cancer types. P-values are highlighted for significant cases only. **(E)** Correlations between EXTEND scores and gene expression of ten oncogenic signaling pathways across 31 cancer types. Significant correlations (FDR < 0.05) are shown in either red (positive correlation) or blue circles (negative correlation). Frequency of each gene’s positive and negative correlation patterns across different cancer types is summarized on the right.

Using seven cancer types where each of the four stages had at least 10 samples, we found in four of seven tested cancer types (THCA, KIRC, KIRP, LUAD), EXTEND scores increased in high-stage tumors, suggesting higher telomerase activity in advanced stage tumors consistent with previous reports^58–61^. In contrast, stomach adenocarcinoma (STAD) and colorectal adenocarcinoma (CRC) showed the highest scores in stage I tumors (**Fig. 3B**). In melanoma, metastatic cancers exhibited higher scores than primary and regionally invasive tumors (**Fig. 3C**).

Given the correlation between EXTEND score and tumor stage, we further tested its association with tumor molecular subtypes. EXTEND scores were significantly different in at least one subtype classification in all tested cancer types (**Supplementary Fig. 12**), suggesting telomerase activity may be an important factor underlying the molecular heterogeneity of cancer.

We also examined associations between telomerase activity and patient prognosis. We categorized EXTEND scores into low and high tumor groups based on the median and performed univariate and multivariate survival analyses. Five cancer types (ACC, KICH, KIRP, LUAD and SARC) exhibited worse overall survival in the high score group, whereas stomach adenocarcinoma (STAD) and thymoma (THYM) exhibited the opposite pattern (**Fig. 3D**). However, only ACC and STAD remained significant in the multivariate analysis when controlling for tumor stage and patient age at diagnosis, likely due to the association between telomerase activity and tumor stage.

Next, we correlated EXTEND scores with the ten oncogenic signaling pathways recently curated by PanCan Atlas to identify potential regulators of telomerase^62^. Cell cycle, p53, Myc, and receptor tyrosine kinase (RTK) pathways were largely positively correlated with EXTEND scores, whereas the TGF beta and Wnt pathways were negatively correlated (**Fig. 3E**). The positive correlation between cell cycle genes and EXTEND corroborates the observation that telomere extension by telomerase occurs during cell cycle^42,63^. Myc, Wnt, and TGF beta pathways, specifically c-myc, beta-catenin, and *TGFBR2*, directly regulate telomerase^64–68^. Expression of *PDGFRA*, a marker of mesenchymal cells, was negatively correlated with EXTEND scores in 21 of 31 cancer types. This was in stark contrast to other genes of the RTK pathway, suggesting possible suppression of telomerase activity in cells of mesenchymal origins.

### Telomerase activity and cancer stemness are positively correlated

Increasing evidence suggests cancers exhibit stem-cell like characteristics, though it is still controversial if cancer stemness reflects the presence of cancer stem cells or stem cell-associated programs^69^. Since active telomerase is a feature of stem cells, we examined the association between cancer stemness and telomerase activity.

We obtained stemness measurements of TCGA tumors from a recent pan-cancer analysis^70^. This cancer stemness index was calculated from an expression signature of 12,945 genes independently derived by comparing embryonic stem cells and differentiated progenitor cells. We found cancer stemness and EXTEND scores were highly correlated at the cancer type level (Rho=0.85, P=2e-9) (**Fig. 4A**). This significant correlation remained within each cancer type (FDR<0.05), suggesting it was cancer lineage independent. Nine of the 13 EXTEND signature genes overlapped with the stemness signature. However, removing these nine genes from the stemness signature virtually had no impact on the resulting stemness scores as they only constituted less than 0.1% of the stemness signature. To further validate the correlation between stemness scores and EXTEND, we performed a permutation analysis by randomly shuffling gene labels of the expression data. We then calculated empirical p values by comparing observed correlations with those generated from the random permutation; 27 of 31 cancer types remained highly significant (empirical p values < 0.05) (**Supplementary Table 5**).

**Fig. 4.**
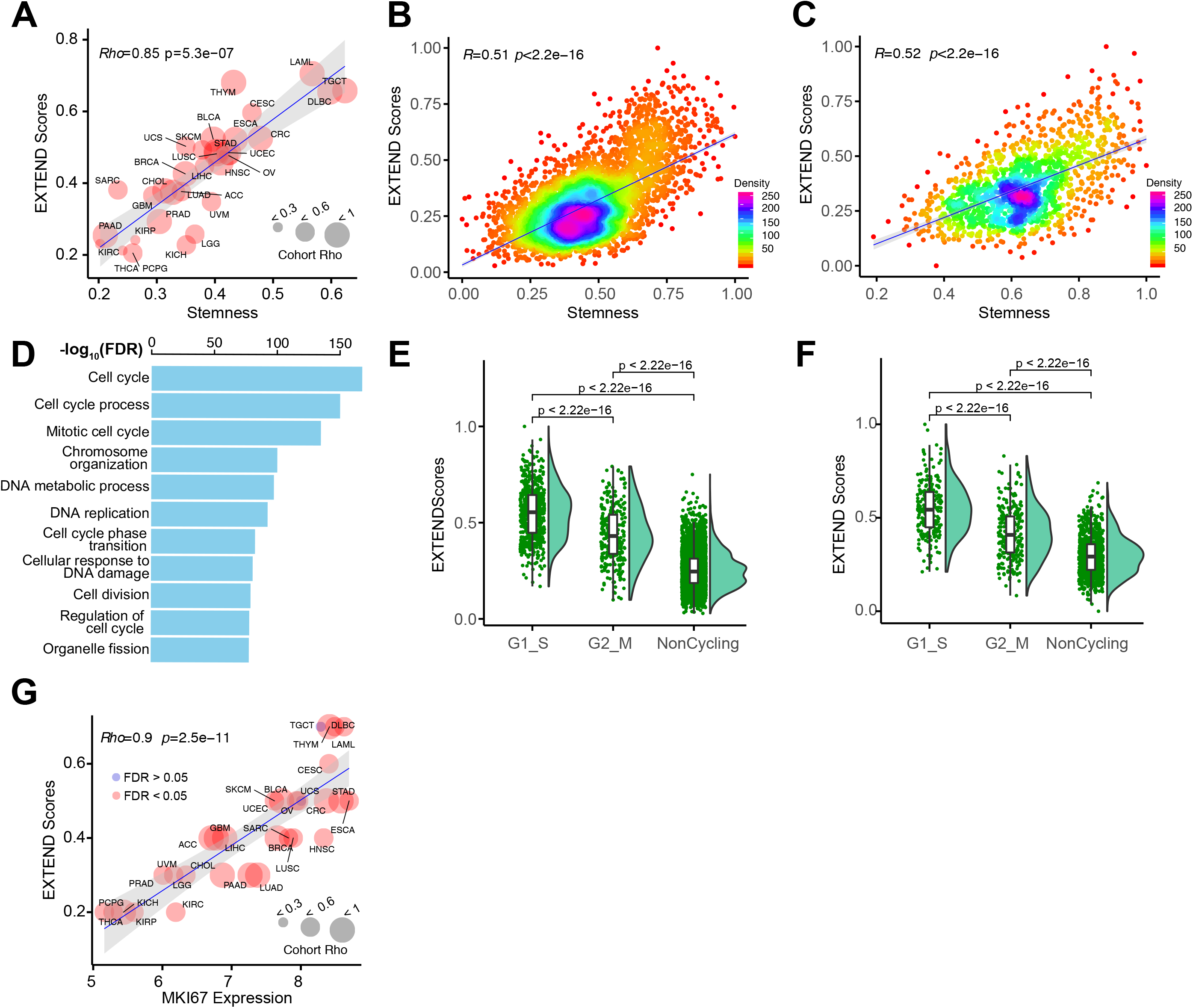
Association between EXTEND score, cancer stemness and proliferation. (**A**) Correlation between telomerase activity and cancer stemness across TCGA cohorts. Node size is proportional to correlation coefficients. In all cancer types, tumor stemness and telomerase activity is significantly correlated (FDR<0.05). (**B-C**) Correlation between EXTEND score and cancer stemness at single cell level in (**B**) glioblastoma and (**C**) head and neck cancer. Each dot represents one cell. (**D**) Top ten pathways enriched in the high stemness, high telomerase cells from head and neck cancer. (**E-F**) EXTEND scores of cycling cells (G1-S phase and G2-M phase) and non-cycling cells in (**E**) glioblastoma (**F**) head and neck cancer. (**G**) Correlation between EXTEND scores and proliferation marker MKI67 expression across 31 cancer types. This correlation is significant in all but 2 cancer types (FDR<0.05).

We next asked if these tumor and tissue level correlations reflected tumor cell behavior. We calculated cancer stemness using single-cell RNAseq data from glioblastoma^71^, head and neck cancer^72^, and medulloblastoma samples^73^. Though none of the datasets had the sensitivity to detect *TERT* or *TERC* expression due to the low input materials from single cells, EXTEND successfully estimated telomerase activity scores for each cell. We again observed strong positive correlations between single cell EXTEND scores and cancer stemness (P < 2.2e−16) (**Figs. 4B, 4C** and **Supplementary Fig. 13A**), suggesting the associations between the two concepts are inherent to cancer cells.

To characterize these high stemness, high telomerase cells, we identified genes and their corresponding pathways differentially expressed in these cells (**Supplementary Table 6, Supplementary Fig. 14**). Cell cycle, DNA replication, and repair pathways were significantly enriched among all three cancer types (**Fig. 4D**), suggesting these were cycling cells. To further verify this finding, we divided cells into G1_S, G2_M, and non-cycling based on recently published markers^71^. Cells in G1_S phase exhibited significantly higher EXTEND scores than G2_M cells and non-cycling cells (**Figs. 4E, 4F and Supplementary Fig 13B**). Taken together, these data support a model that a group of high stemness, high telomerase cells drive tumor growth.

This model also predicts that telomerase activity is not only a marker for tumor stemness but also a marker for tumor proliferation. Although previously documented^74,75^, the relationship between telomerase activity and tumor proliferation has not been quantitatively measured. Using KI67, a cell proliferation marker, we observed a linear positive correlation between tumor proliferation and EXTEND score on cancer type level (Rho=0.9, P=2.5e-11) and in most tumor cohorts (FDR<0.05). Exceptions were two tumor types that originate from reproductive organs, uterine carcinosarcoma (UCS) and testicular germ cell tumors (TGCT) (**Fig. 4G**).

## Discussion

*TERT* has long been recognized as a pivotal determinant of telomerase enzymatic activity. However, it has been increasingly clear that *TERT* is involved in telomere independent functions and its expression has limitations in predicting telomerase activity^23,32–34^. Furthermore, *TERT* has a high GC content (58% vs genome-wide average 41% in human), making it hard to capture in sequencing even for bulk samples. This issue is exacerbated by the low expression of *TERT*, which was estimated to be between 1-40 copies per cell in cancer cells^34^. Indeed, only 73% of TCGA tumors detected *TERT* expression^76^, a fraction significantly lower than the TRAP assay-based estimate of telomerase positive tumors (~90%)^22^. In single cell RNA sequencing studies, it is typical that none of *TERT* reads are detected in any cells. Thus, *TERT* expression is inadequate, both biologically and technically, for guiding a systematic analysis of telomerase activity.

We here developed EXTEND to predict telomerase activity based on gene expression data. Using cancer cell lines of various tissues of origin, we show that EXTEND outperformed *TERT* expression in predicting telomerase activity. Analyses of GTEx and embryonic samples further supported this conclusion. Interestingly, brain tissues exhibited detectable *TERT* expression in the GTEx data but low EXTEND scores, likely due to *TERT* alternative splicing leading to isoforms that do not encode catalytically active proteins. The brain was reported to have an extreme transcriptome diversity due to alternative splicing compared with other tissues^77^, thus the excessive alternative splicing of *TERT* is not surprising and unlikely specific to *TERT*. Moreover, EXTEND accurately demonstrated the point where telomerase activity is diminished during fetal heart development. These data substantiate EXTEND’s validity and distinguish it from *TERT*. An important reason for EXTEND’s robust performance is the eleven marker genes that have no reported functional associations with the telomerase. These genes not only reduce the impact of *TERT* expression on the final scores, but also provide an avenue for estimating telomerase activity even when *TERT* or *TERC* are beyond detection sensitivity in bulk or single cell samples.

Applying EXTEND to >9,000 tumors and 700 normal samples confirmed many previously known associations but also revealed important new insights. Telomerase activity, as quantified by EXTEND, is continuous rather than dichotomous. Variations in telomerase activity across cancer types can be partially explained by *TERT* expression (R=0.54, P=0.003). Strikingly, cancer stemness, a measure reflecting the overall similarity in self-renewal between cancer cells and embryonic stem cells, correlates significantly better than *TERT* expression with telomerase activity (Rho=0.85, P=2e-9). This correlation coefficient indicates 72% of the variation in telomerase activity across cancer types can be explained by their differences in cancer stemness. Single cell analysis confirmed the tight correlation between active telomerase and cancer stemness program, and further revealed the high stemness high telomerase cells were cycling cells. Future studies will be needed to elucidate the identifies of these cells, and whether inhibiting their telomerase activity, or targeting the stemness program, or both can effectively eliminate these cells.

The lack of correlation between telomerase activity and telomere lengths was surprising but could be explained by several reasons. First of all, telomere lengths are determined by the counteracting effects of attrition during cell division and extension by telomerase. Second, telomere lengths used in our analysis were estimated based on abundances of telomeric repeats from sequencing data^76^. At best, short read sequencing data can only estimate the average telomere length of a sample. However, telomerase preferentially acts on the shortest telomeres^5^. Furthermore, a recent study suggests telomeric repeats can frequently insert in non-telomeric regions of the genome ^78^. Thus, overall telomeric repeat counts may not precisely measure a tumor’s telomere lengths.

In summary, our study demonstrates the feasibility of digitalizing telomerase enzymatic activity, a pathway fundamental to cancer cell and stem cell survival. Our analysis establishes quantitative associations between telomerase activity and cancer stemness and proliferation.

## Acknowledgement

This work was supported by CPRIT (RR170055 to S.Z.) and MD Anderson Cancer Center Brain Cancer SPORE Program Career Enhancement Award (S.Z.). We thank Dr. Zhu Wang (UT Health San Antonio) for discussions on statistical issues and Karen Klein for editorial assistance. The results shown here are in whole or part based upon data generated by the TCGA Research Network. The Genotype-Tissue Expression (GTEx) Project was supported by the Common Fund of the Office of the Director of the National Institutes of Health, and by NCI, NHGRI, NHLBI, NIDA, NIMH, and NINDS. The data used for the analyses described in this manuscript were obtained from the GTEx Portal.

## Supplementary Figure Legends

**Supplementary Fig. 1.** Schematic of EXpression-based Telomerase Enzymatic activity Detection (EXTEND) tool workflow.

**Supplementary Fig. 2.**(**a**) Frequencies of ATRX/DAXX alteration and TERT promoter mutation across TCGA cancer cohorts. X-axis indicates sample size. Each bar represents one cancer type from TCGA. (**b**) TERT promoter mutation and ATRX mutations are strongly mutually exclusive in LGG.

**Supplementary Fig. 3.** Contribution of signature genes towards EXTEND score evaluated using Cancer Cell Line Encyclopedia (CCLE) data. Each dot represents one cell line. TERT/TERC contributes more than other genes in the signature but overall the contribution is less than 20%.

**Supplementary Fig. 4.** Spearman correlation between PolyA enrichment protocol-based prediction and ribosomal depletion protocol-based prediction. Data from GSE51783 was used for this analysis. X-axis represents PolyA enrichment protocol-based scores, whereas y-axis represents ribosomal depletion protocol-based scores.

**Supplementary Fig. 5.** Spearman correlation of telomerase activity (measured by TRAP assay) with **(a)** EXTEND scores and **(b)** TERT expression across 28 glioma cell lines. Each dot represents a cell line. X-axis in both panels denotes telomerase activity from TRAP assays. Y-axis is EXTEND score and TERT expression, respectively. GSC5-22 and GSC818 are two ALT lines.

**Supplementary Fig. 6.** Spearman correlation of telomerase activity (obtained through enzymatic assay) with **(a)** EXTEND scores and **(b)** TERT expression across 11 bladder cancer cell lines. Correlation coefficient (Rho = 0.72, P-value = 0.013) of EXTEND scores versus telomerase activity significantly outperforms TERT expression (Rho= 0.55, P-value = 0.083).

**Supplementary Fig. 7.** Spearman correlation of telomerase activity (obtained through digital droplet-based TRAP assay) with **(a)** EXTEND scores and **(b)** TERT expression across 15 lung cancer cell lines. Correlation coefficient (Rho = 0.65, P-value = 0.01) of EXTEND scores versus telomerase activity significantly outperforms TERT expression (Rho= 0.48, P-value = 0.07).

**Supplementary Fig. 8.** Distribution of EXTEND scores across human tissues. Each group represents one tissue type from GTEx. TERT expression is denoted by the left y axis, whereas EXTEND score is denoted by the right y axis.

**Supplementary Fig. 9.** A screenshot from UCSC Xena shows the percentages of TERT isoforms in brain (purple) versus testis (cyan) across GTEx samples. Most of TERT expressed in the brain are short spliced forms (second last), whereas the proportion of full-length isoform is close to zero. In contrast, about 25-30% of TERT expressed in the testis are full-length isoforms.

**Supplementary Fig. 10.** Correlation between EXTEND and POT1 across pan-cancer (left). No correlation was observed between POT1 and TERT expression (right). For this analysis, EXTEND scores and gene expression were z-score transformed in each cancer cohort to remove tissue effect.

**Supplementary Fig. 11.** Correlation between telomere lengths (obtained from Barthel et al.,2017) and EXTEND scores across 31 cancer types (TCGA) for WGS and WXS platforms. Numbers in each cell (of heatmap) represent spearman correlation coefficient while the color represents adjusted p value (multiple testing via Bonferroni correction).

**Supplementary Fig. 12.** EXTEND scores across various subtypes in cancer cohorts (TCGA data). P-values were calculated using two-sided t-test.

**Supplementary Fig. 13.** Correlation between telomerase activity and cancer stemness in medulloblastoma. **(a)** Scatter plot representing correlation (spearman rank method) of EXTEND scores with stemness index for single cell data from Medulloblastoma (Hovestadt et al., 2019). **(b)** Violin plot representing distribution of EXTEND scores across cycling cells (in G1-S and G2-M phase) and noncycling cells in Medulloblastoma data. P-values were calculated using two-sided t-test.

**Supplementary Fig. 14.** Distribution of EXTEND scores and cancer stemness scores. Their distributions consistently show multi-modes. Based on these distributions, we set EXTEND score to 0.5 as the cutoff to select for high stemness, high telomerase activity cells in the three single cell datasets (accounting for 10% of glioblastoma cells, 17% of head and neck cells, and 14% of medullo cells). Linear regression gives a slope of 0.67 for medulloblastoma, and 0.6 for both glioblastoma and head and neck cancer when considering EXTEND score as the dependent variable. Intercept is close to 0 for all three in linear regression.

## Supplementary Table Legends

**Supplementary Table 1:** Telomerase Signature Genes information.

**Supplementary Table 2:** Pathway enrichment of telomerase signature genes retrieved through MSigDb.

**Supplementary Table 3:** EXTEND Scores for samples cross Genotype-Tissue Expression (GTEx) data.

**Supplementary Table 4:** EXTEND Scores for 31 cancer types in TCGA Pancan.

**Supplementary Table 5:** EXTEND Scores and Stemness index Correlation based on Permutation Test across TCGA cohort

**Supplementary Table 6:** Pathways enrichment (MSigDb) for Up and downregulated genes based on High and Low EXTEND Scores for 3 single cell data sets (Glioblastoma, Medulloblastoma and Head and Neck cancers)

