## Supplementary Figures for "Inferring Telomerase Enzymatic Activity from Expression Data"

### Slide 1
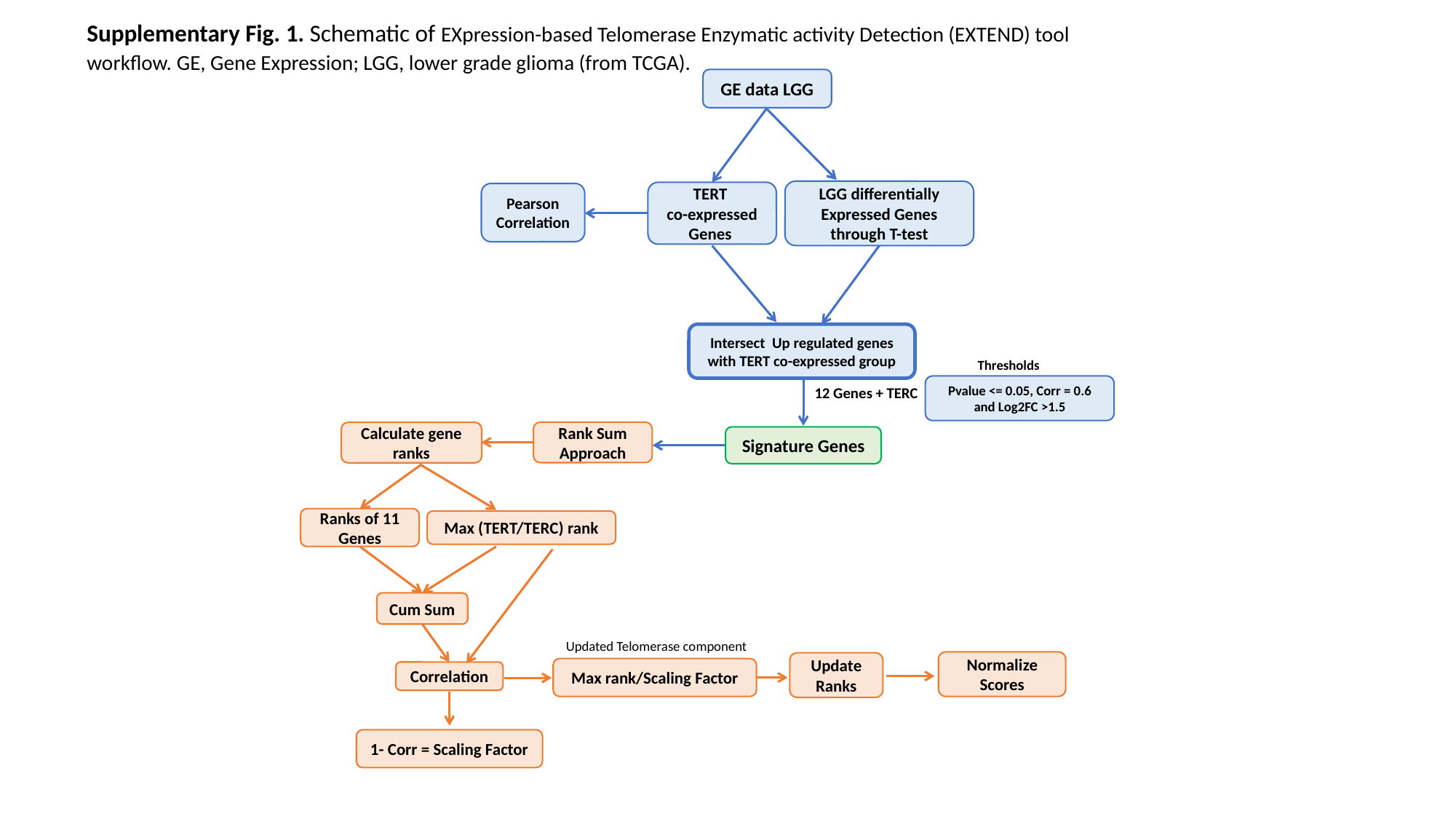

Supplementary Fig. 1. Schematic of EXpression-based Telomerase Enzymatic activity Detection (EXTEND) tool workflow. GE, Gene Expression; LGG, lower grade glioma (from TCGA).
GE data LGG
LGG differentially Expressed Genes through T-test
TERT
co-expressed Genes
Pearson Correlation
Intersect Up regulated genes with TERT co-expressed group
Thresholds
Pvalue <= 0.05, Corr = 0.6 and Log2FC >1.5
12 Genes + TERC
Rank Sum Approach
Calculate gene ranks
Signature Genes
Ranks of 11 Genes
Max (TERT/TERC) rank
Cum Sum
Updated Telomerase component
Normalize Scores
Update Ranks
Max rank/Scaling Factor
Correlation
1- Corr = Scaling Factor

### Slide 2
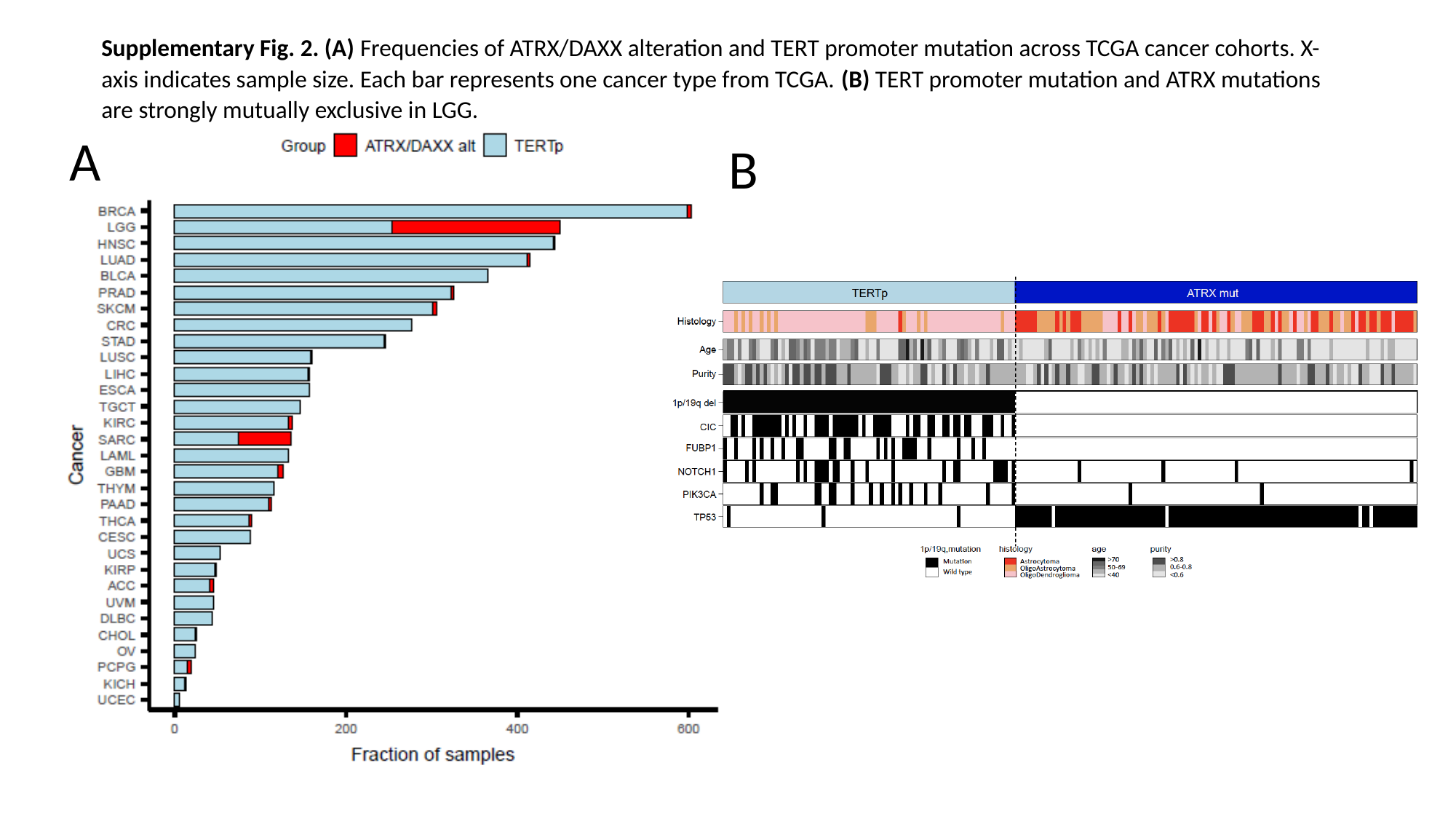

Supplementary Fig. 2. (A) Frequencies of ATRX/DAXX alteration and TERT promoter mutation across TCGA cancer cohorts. X-axis indicates sample size. Each bar represents one cancer type from TCGA. (B) TERT promoter mutation and ATRX mutations are strongly mutually exclusive in LGG.
A
B

### Slide 3
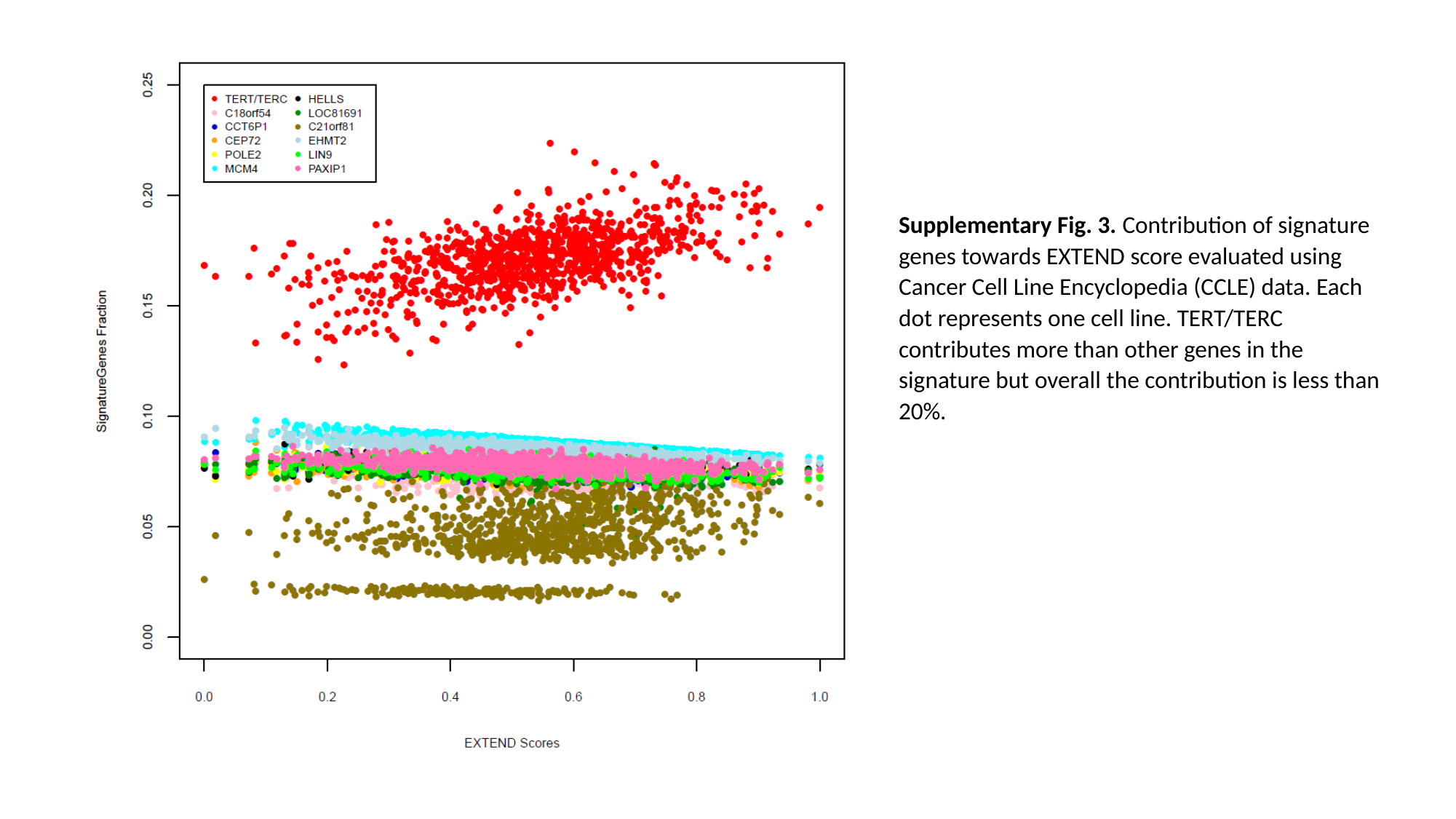

Supplementary Fig. 3. Contribution of signature genes towards EXTEND score evaluated using Cancer Cell Line Encyclopedia (CCLE) data. Each dot represents one cell line. TERT/TERC contributes more than other genes in the signature but overall the contribution is less than 20%.

### Slide 4
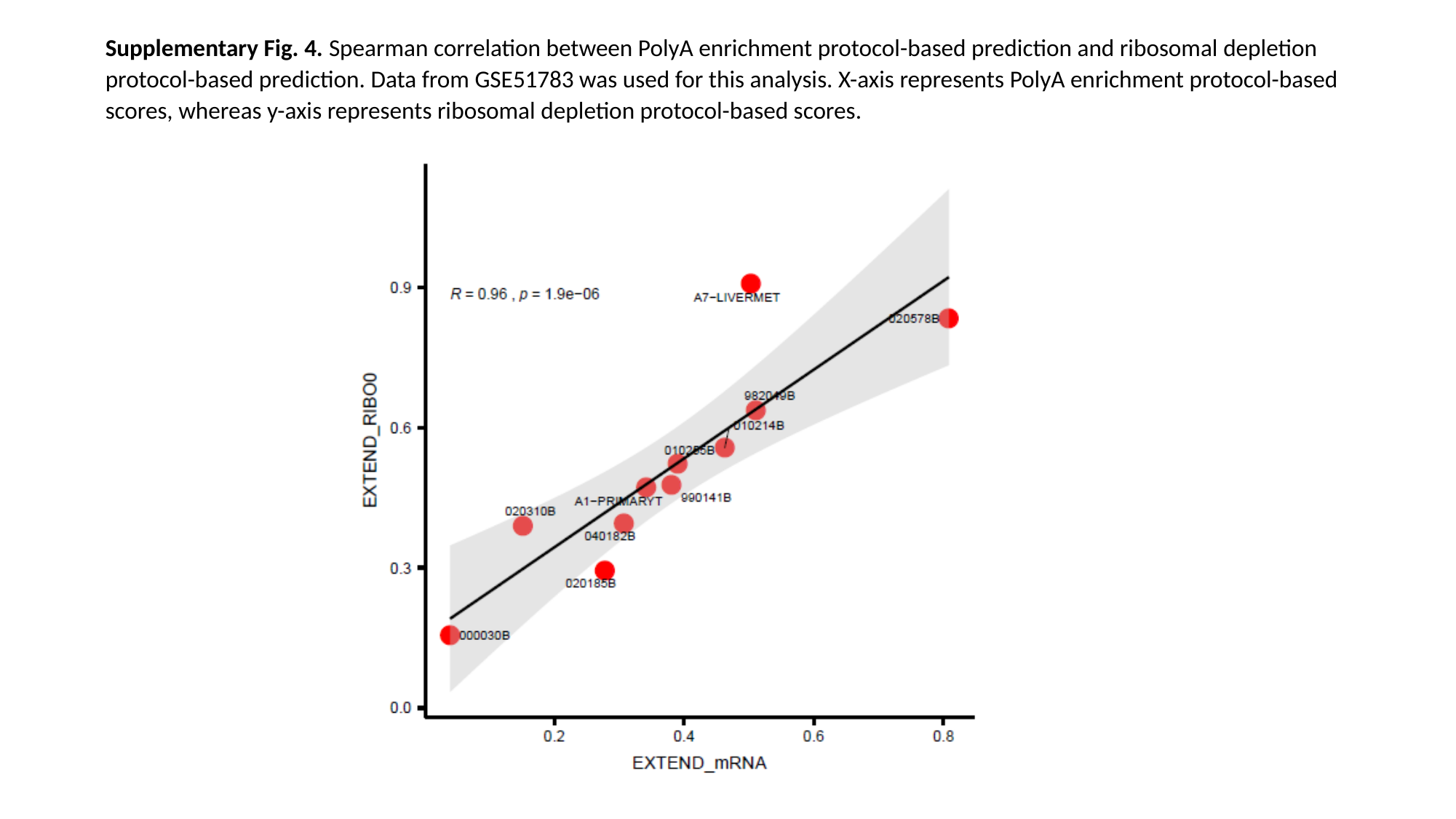

Supplementary Fig. 4. Spearman correlation between PolyA enrichment protocol-based prediction and ribosomal depletion protocol-based prediction. Data from GSE51783 was used for this analysis. X-axis represents PolyA enrichment protocol-based scores, whereas y-axis represents ribosomal depletion protocol-based scores.

### Slide 5
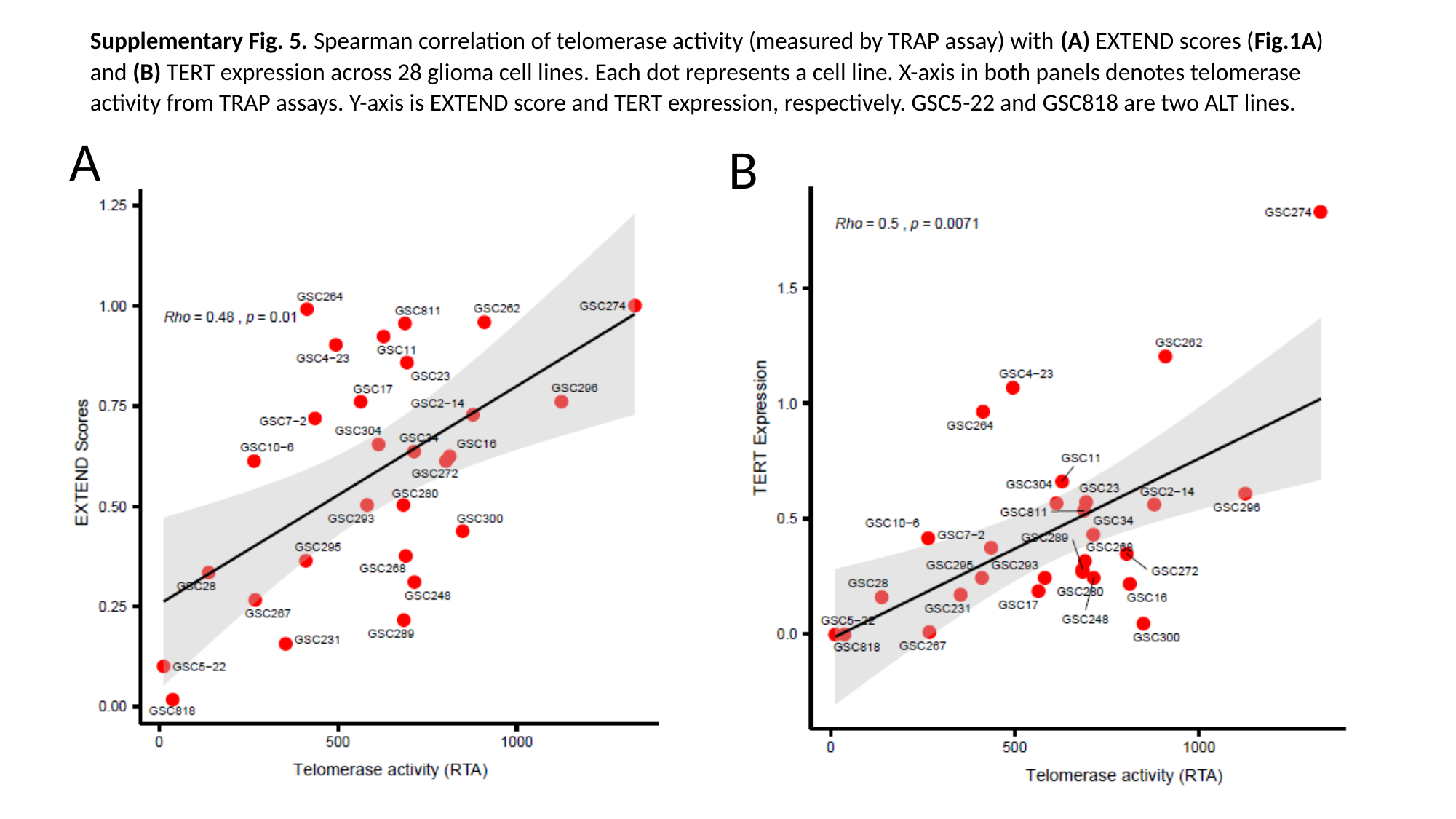

Supplementary Fig. 5. Spearman correlation of telomerase activity (measured by TRAP assay) with (A) EXTEND scores (Fig.1A) and (B) TERT expression across 28 glioma cell lines. Each dot represents a cell line. X-axis in both panels denotes telomerase activity from TRAP assays. Y-axis is EXTEND score and TERT expression, respectively. GSC5-22 and GSC818 are two ALT lines.
A
B

### Slide 6
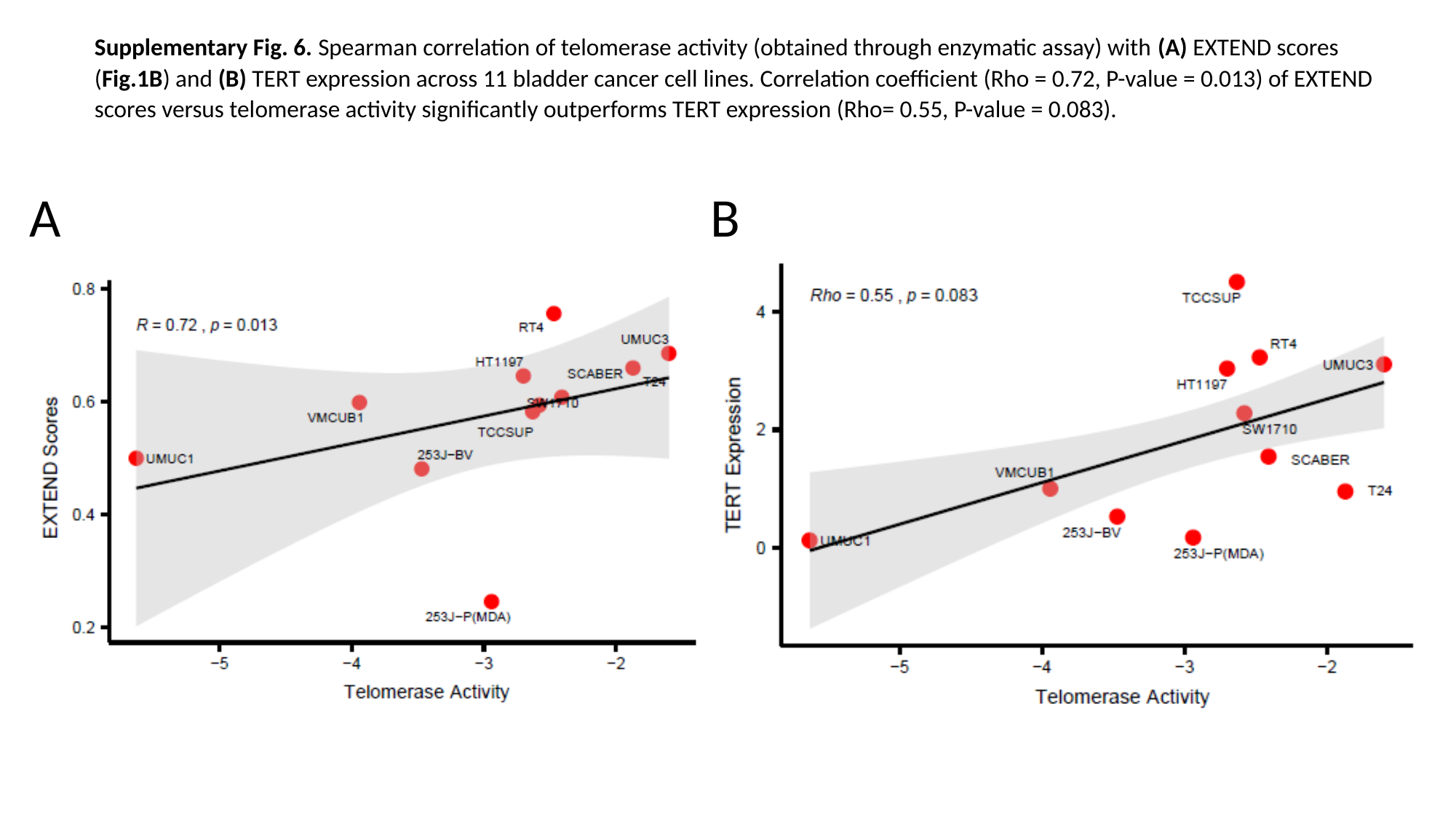

Supplementary Fig. 6. Spearman correlation of telomerase activity (obtained through enzymatic assay) with (A) EXTEND scores (Fig.1B) and (B) TERT expression across 11 bladder cancer cell lines. Correlation coefficient (Rho = 0.72, P-value = 0.013) of EXTEND scores versus telomerase activity significantly outperforms TERT expression (Rho= 0.55, P-value = 0.083).
A
B

### Slide 7
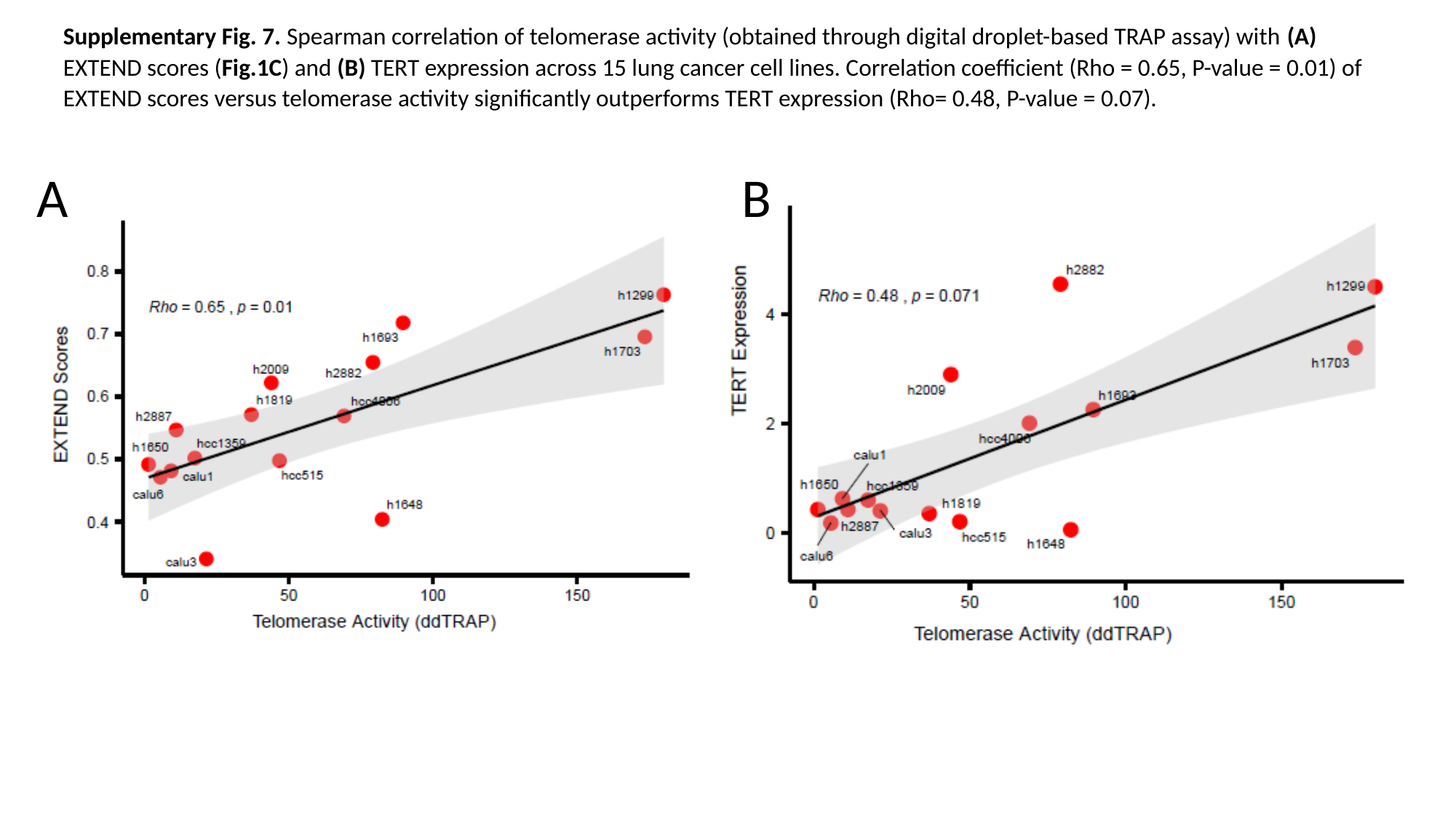

Supplementary Fig. 7. Spearman correlation of telomerase activity (obtained through digital droplet-based TRAP assay) with (A) EXTEND scores (Fig.1C) and (B) TERT expression across 15 lung cancer cell lines. Correlation coefficient (Rho = 0.65, P-value = 0.01) of EXTEND scores versus telomerase activity significantly outperforms TERT expression (Rho= 0.48, P-value = 0.07).
A
B

### Slide 8
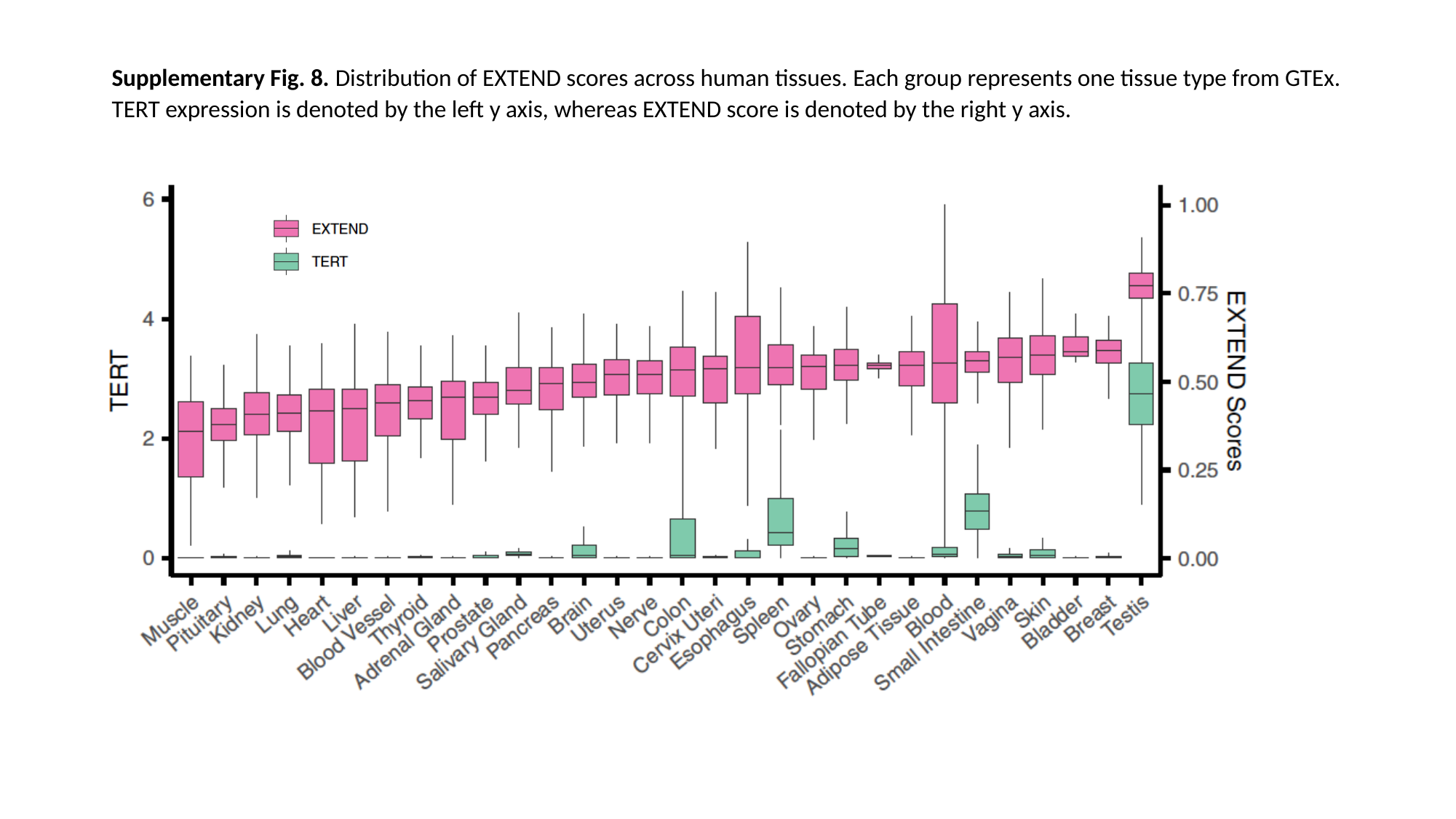

Supplementary Fig. 8. Distribution of EXTEND scores across human tissues. Each group represents one tissue type from GTEx. TERT expression is denoted by the left y axis, whereas EXTEND score is denoted by the right y axis.

### Slide 9
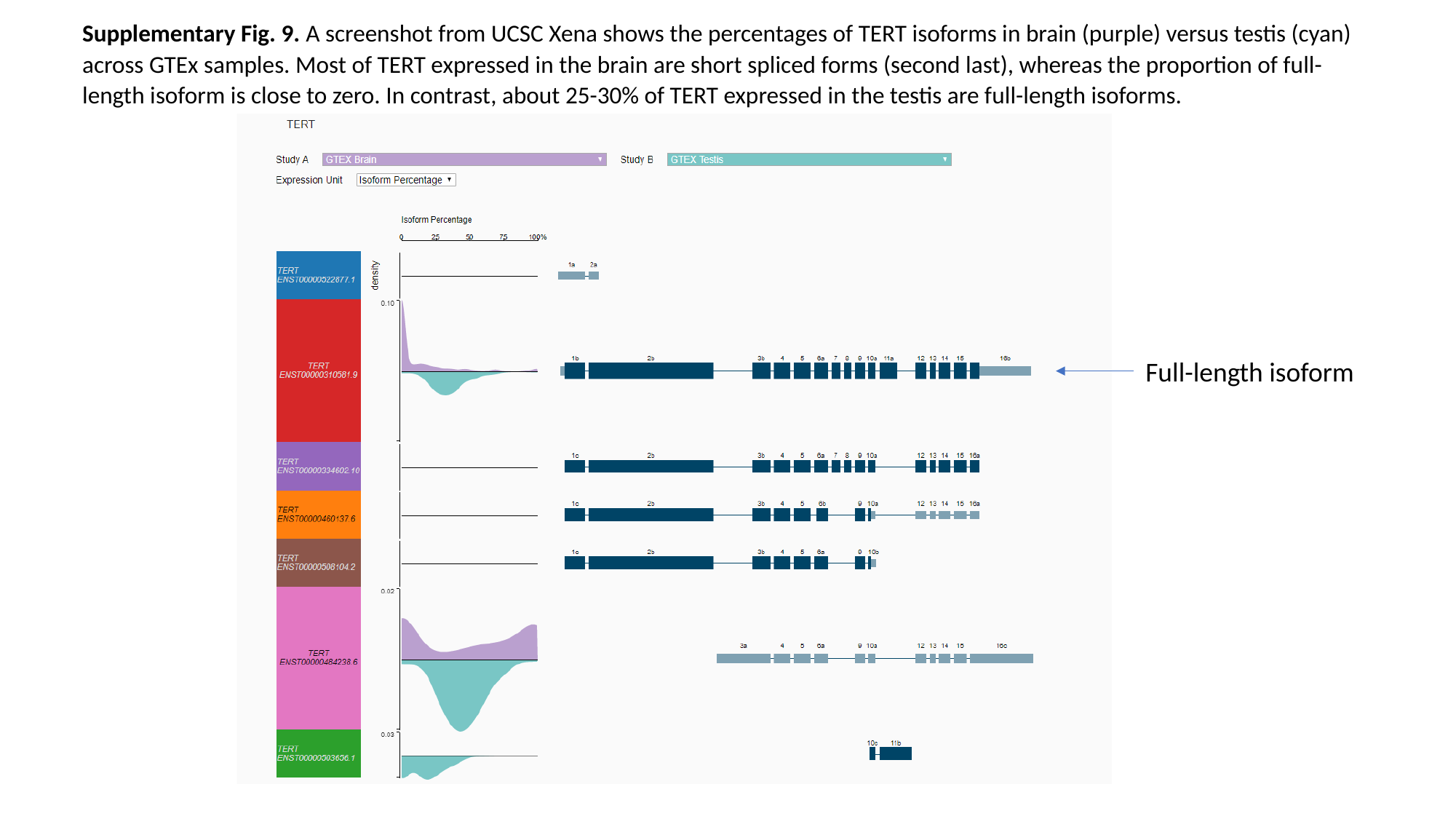

Supplementary Fig. 9. A screenshot from UCSC Xena shows the percentages of TERT isoforms in brain (purple) versus testis (cyan) across GTEx samples. Most of TERT expressed in the brain are short spliced forms (second last), whereas the proportion of full-length isoform is close to zero. In contrast, about 25-30% of TERT expressed in the testis are full-length isoforms.
Full-length isoform

### Slide 10
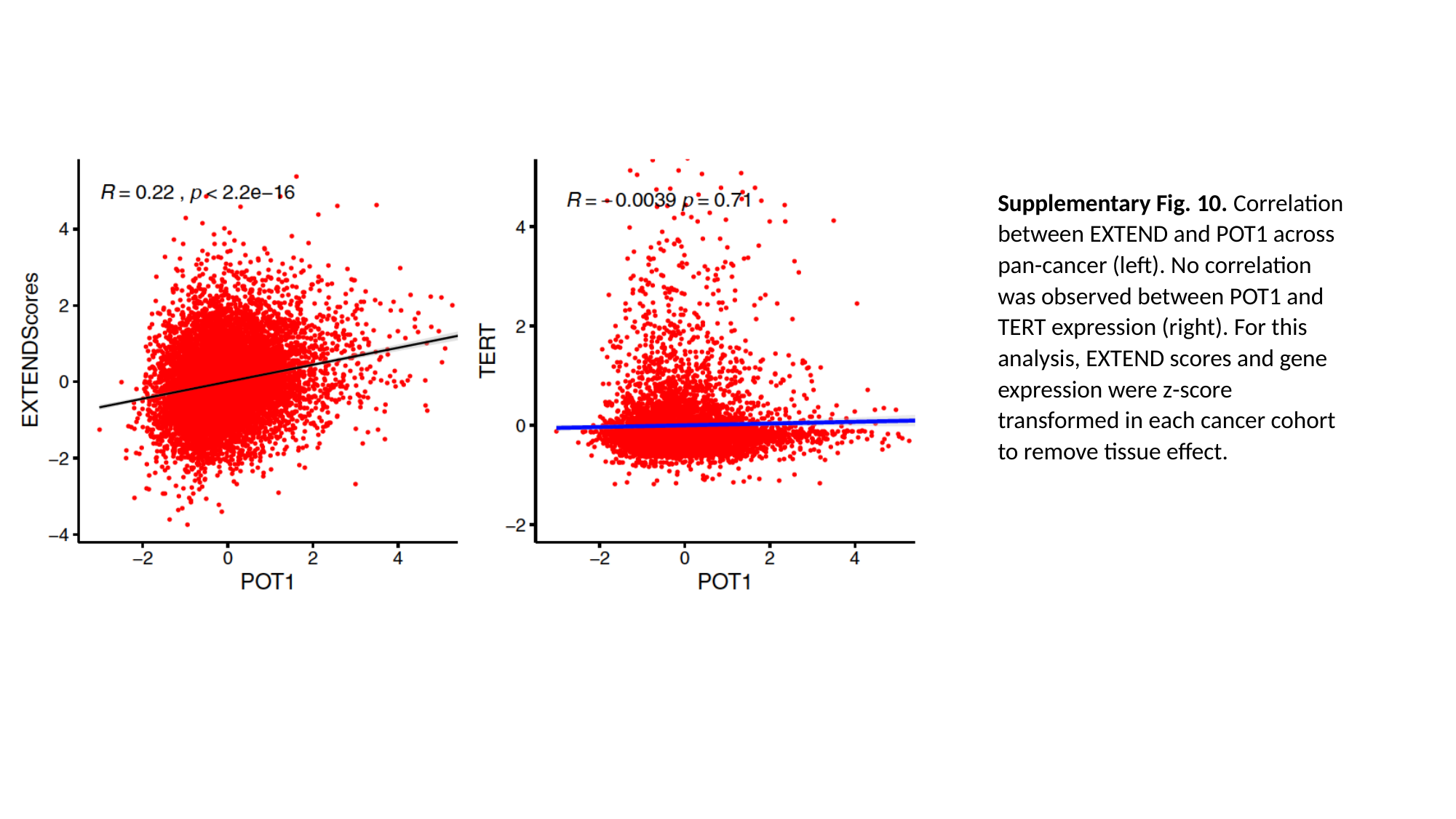

Supplementary Fig. 10. Correlation between EXTEND and POT1 across pan-cancer (left). No correlation was observed between POT1 and TERT expression (right). For this analysis, EXTEND scores and gene expression were z-score transformed in each cancer cohort to remove tissue effect.

### Slide 11
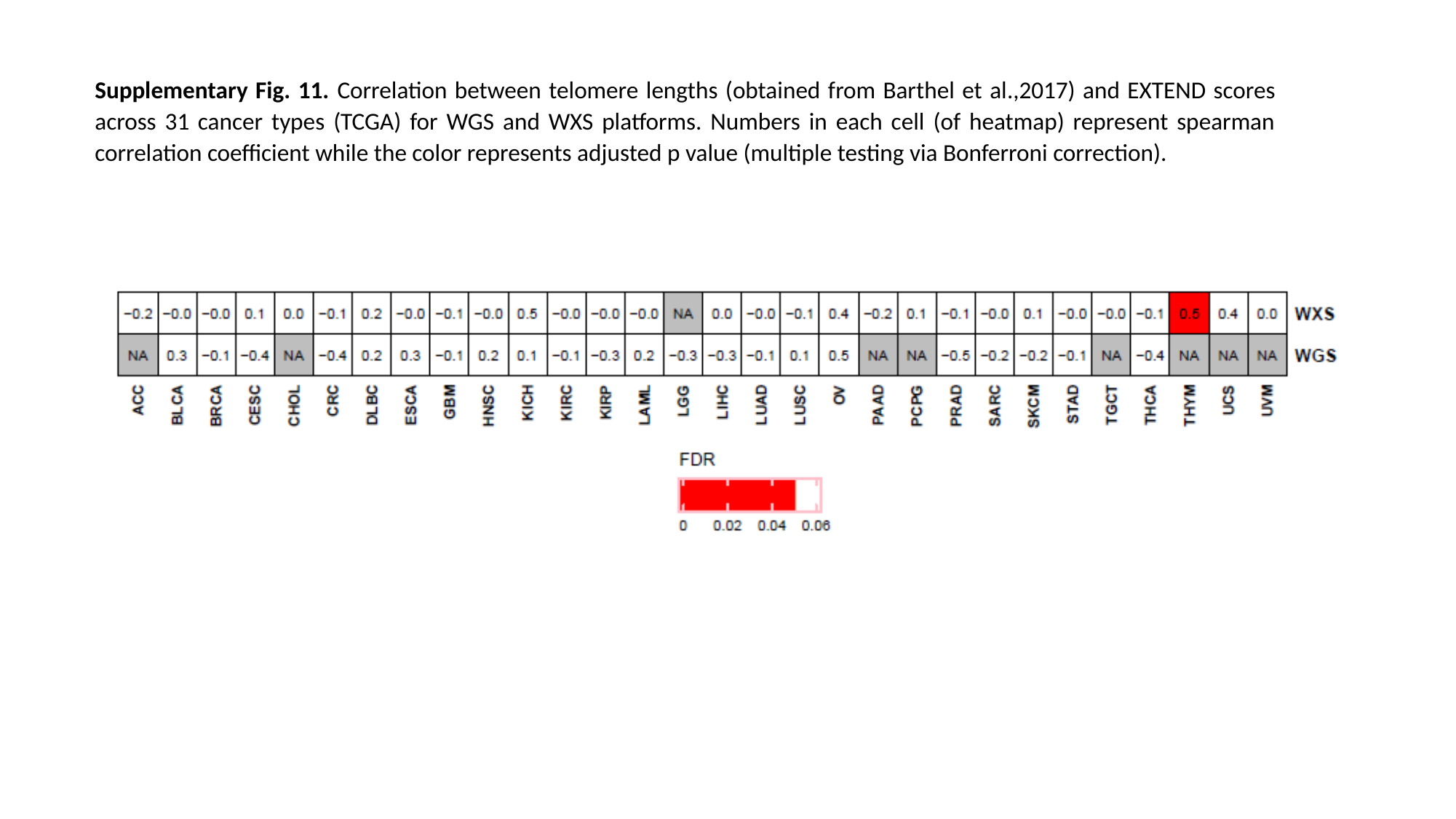

Supplementary Fig. 11. Correlation between telomere lengths (obtained from Barthel et al.,2017) and EXTEND scores across 31 cancer types (TCGA) for WGS and WXS platforms. Numbers in each cell (of heatmap) represent spearman correlation coefficient while the color represents adjusted p value (multiple testing via Bonferroni correction).

### Slide 12
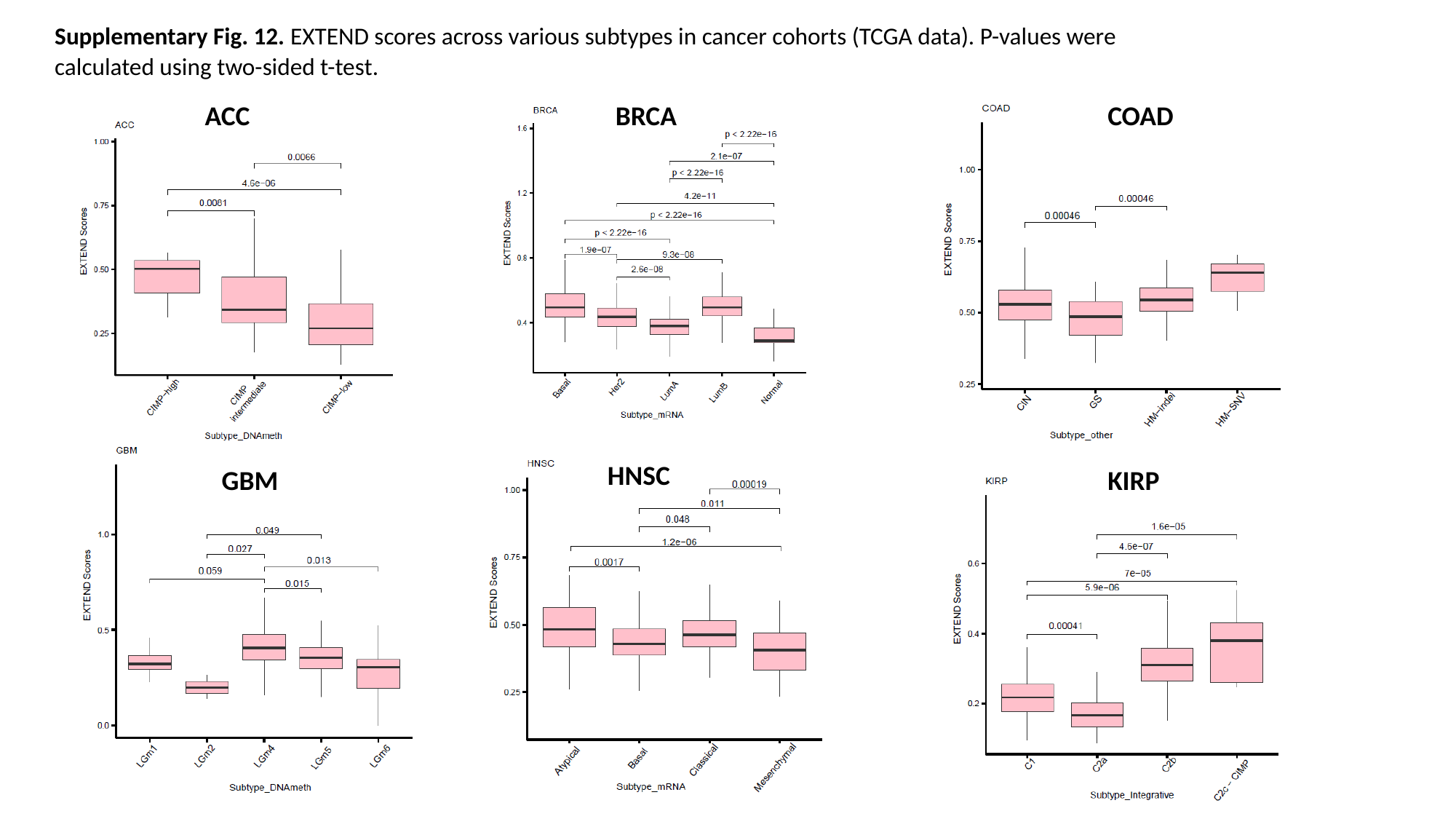

Supplementary Fig. 12. EXTEND scores across various subtypes in cancer cohorts (TCGA data). P-values were calculated using two-sided t-test.
BRCA
COAD
ACC
HNSC
GBM
KIRP

### Slide 13
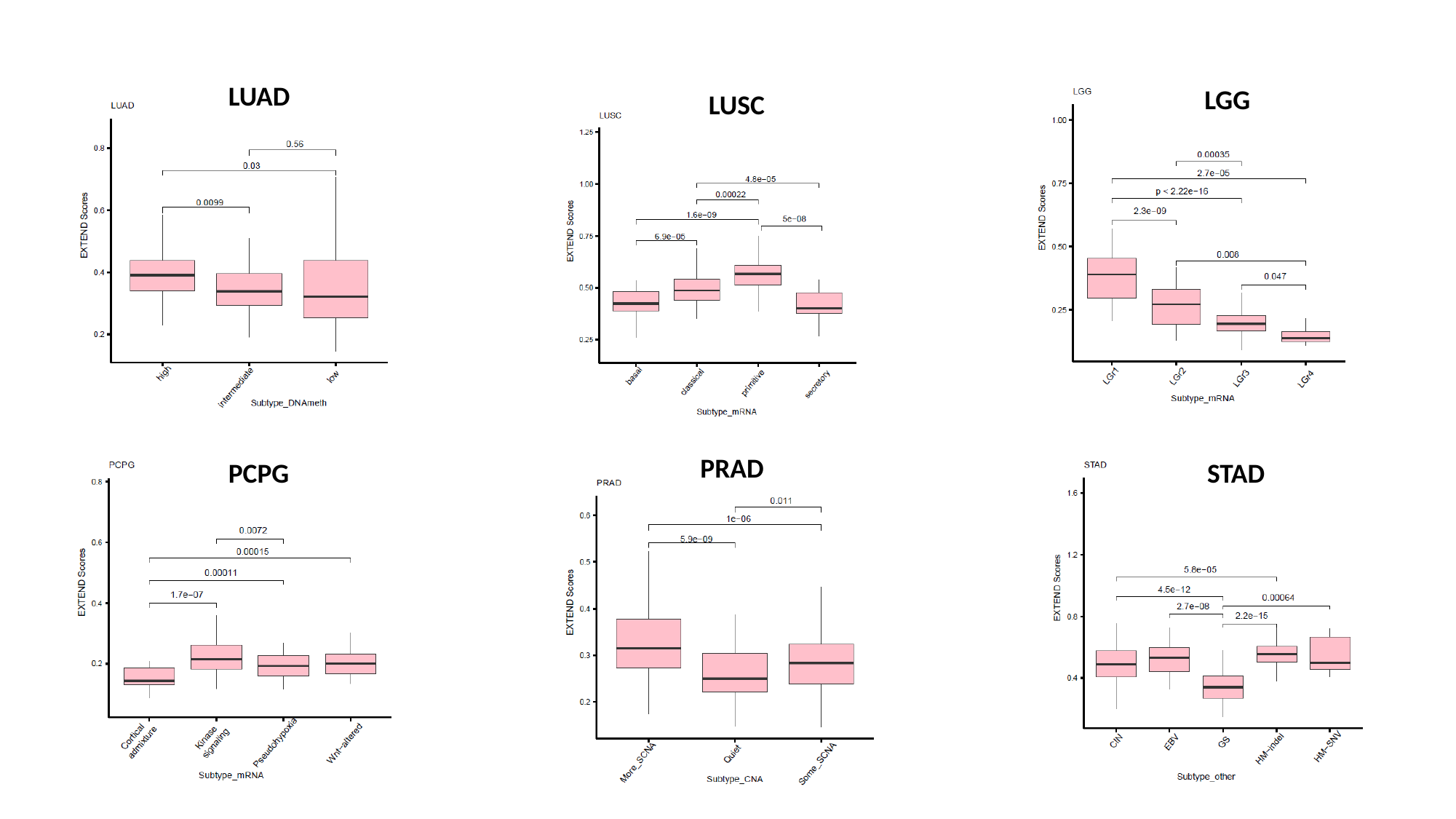

LUAD
LGG
LUSC
PRAD
STAD
PCPG

### Slide 14
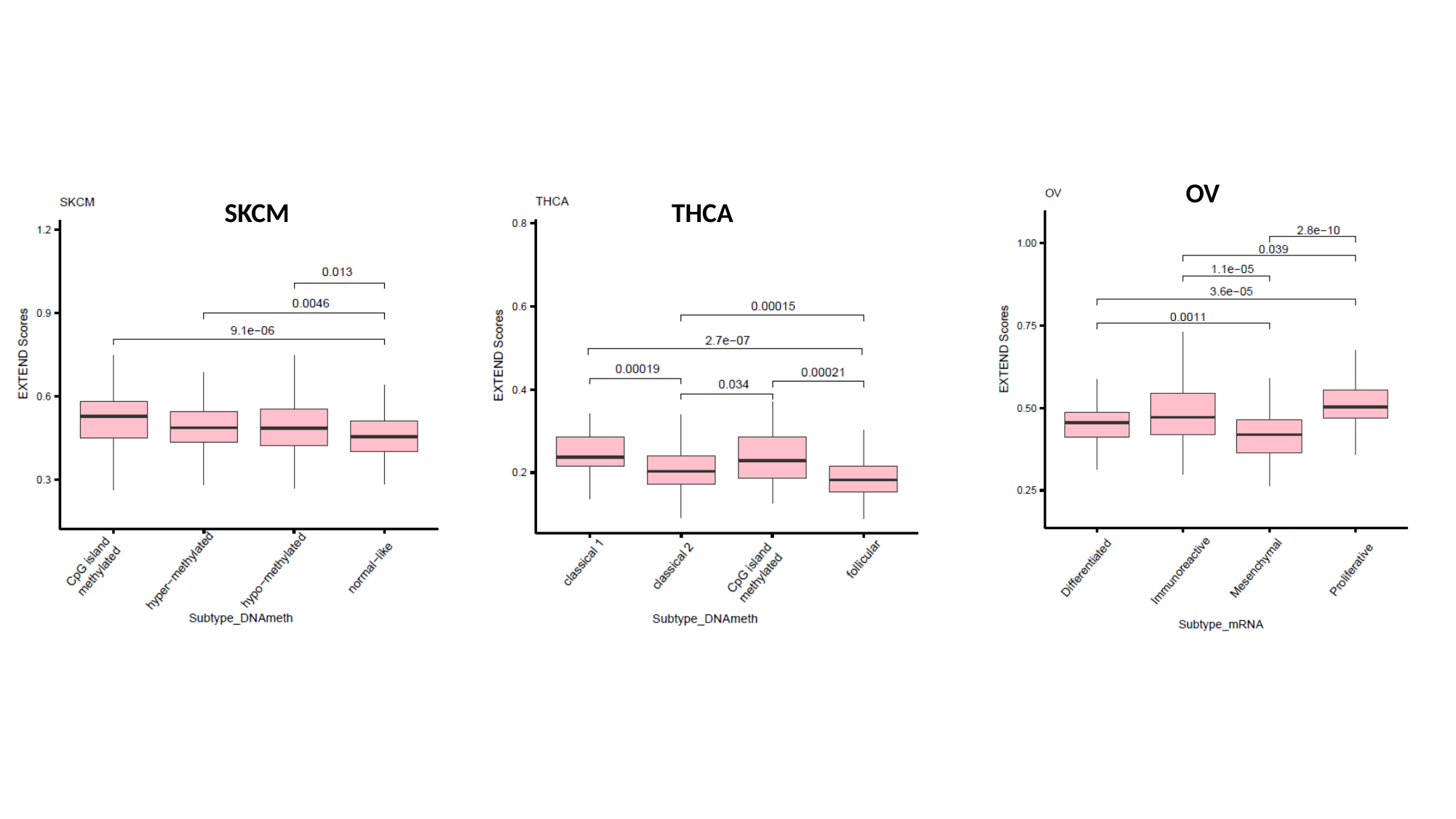

OV
SKCM
THCA

### Slide 15
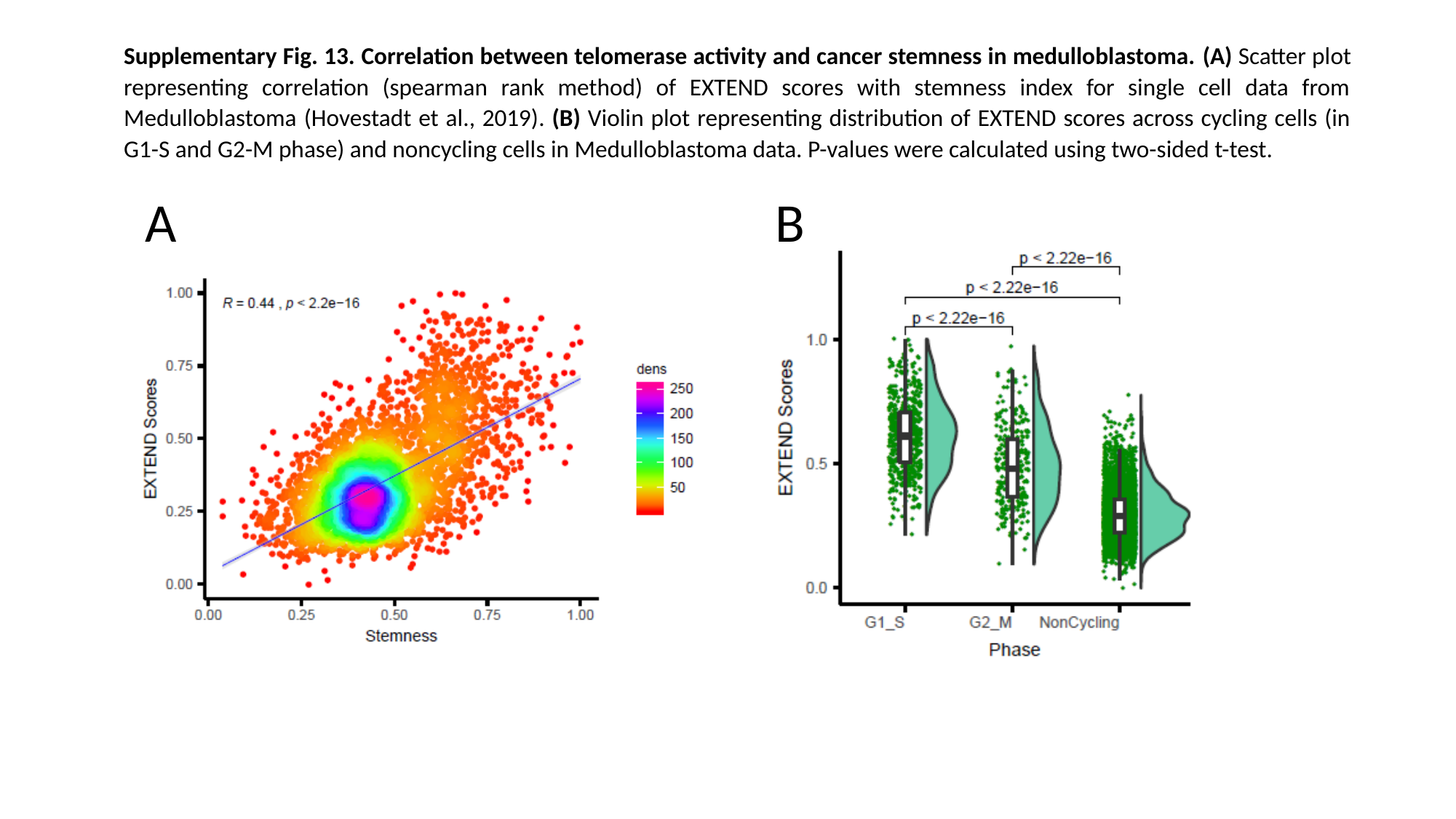

Supplementary Fig. 13. Correlation between telomerase activity and cancer stemness in medulloblastoma. (A) Scatter plot representing correlation (spearman rank method) of EXTEND scores with stemness index for single cell data from Medulloblastoma (Hovestadt et al., 2019). (B) Violin plot representing distribution of EXTEND scores across cycling cells (in G1-S and G2-M phase) and noncycling cells in Medulloblastoma data. P-values were calculated using two-sided t-test.
B
A

### Slide 16
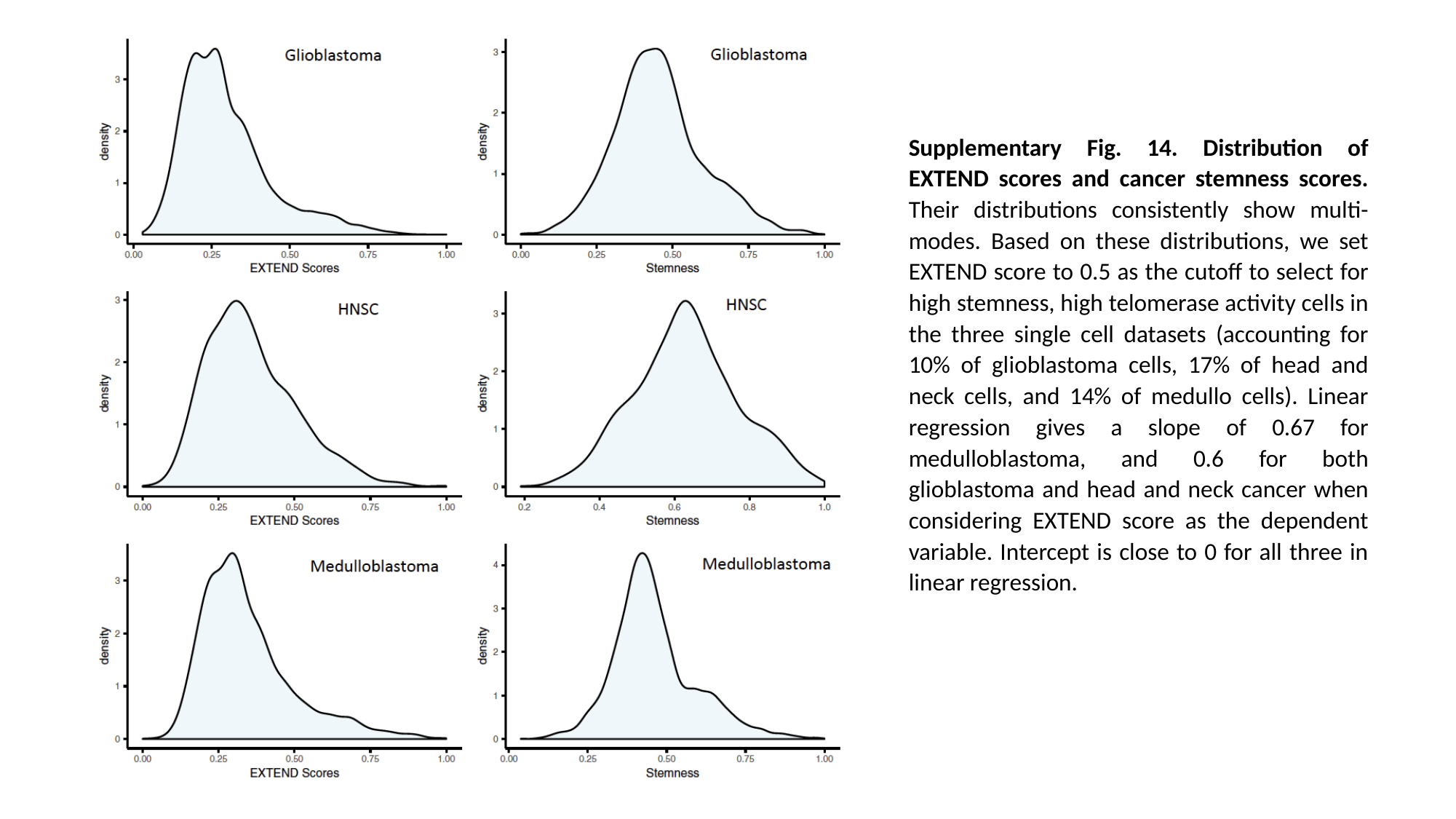

Supplementary Fig. 14. Distribution of EXTEND scores and cancer stemness scores. Their distributions consistently show multi-modes. Based on these distributions, we set EXTEND score to 0.5 as the cutoff to select for high stemness, high telomerase activity cells in the three single cell datasets (accounting for 10% of glioblastoma cells, 17% of head and neck cells, and 14% of medullo cells). Linear regression gives a slope of 0.67 for medulloblastoma, and 0.6 for both glioblastoma and head and neck cancer when considering EXTEND score as the dependent variable. Intercept is close to 0 for all three in linear regression.
